# Inhibiting Runx1 protects heart function after myocardial infarction

**DOI:** 10.1101/2022.02.17.480749

**Authors:** Tamara P Martin, Eilidh A MacDonald, Ashley Bradley, Holly Watson, Priyanka Saxena, Eva A. Rog-Zielinska, Simon Fisher, Ali Ali Mohamed Elbassioni, Ohood Almuzaini, Catriona Booth, Morna Campbell, Pawel Herzyk, Karen Blyth, Colin Nixon, Lorena Zentilin, Colin Berry, Thomas Braun, Mauro Giacca, Martin W McBride, Stuart A Nicklin, Ewan R Cameron, Christopher M Loughrey

**Affiliations:** British Heart Foundation Glasgow Cardiovascular Research Centre, Institute of Cardiovascular & Medical Sciences, University of Glasgow, UK.; Institute for Experimental Cardiovascular Medicine, University Heart Centre Freiburg / Bad Krozingen, Faculty of Medicine, Freiburg, Germany.; Department of Cardiothoracic Surgery, Suez Canal University; Institute of Molecular, Cell and Systems Biology, University of Glasgow, UK.; Glasgow Polyomics, University of Glasgow, Garscube Campus, UK; Institute of Cancer Sciences, University of Glasgow, Glasgow, UK.; Cancer Research UK Beatson Institute, Garscube Estate, Glasgow, UK.; Molecular Medicine Laboratory, International Centre for Genetic Engineering and Biotechnology, Trieste, Italy.; Department of Cardiac Development and Remodelling, Max Planck Institute for Heart and Lung Research, Bad Nauheim, Germany.; School of Cardiovascular Medicine and Sciences, King’s College London British Heart Foundation Centre, London, UK.; School of Veterinary Medicine, University of Glasgow, Garscube Campus, UK.

**Keywords:** Runx1, cardiomyocytes, sarcoplasmic reticulum, calcium handling, myocardial infarction border zone, adverse cardiac remodelling.

## Abstract

Myocardial infarction is a major cause of death worldwide. Effective treatments are required that limit adverse cardiac remodelling and preserve cardiac contractility following myocardial infarction, with the aim of improving patient outcomes and preventing progression to heart failure. The perfused but hypocontractile myocardium bordering a newly created infarct is functionally distinct from the remote surviving myocardium; it is also a major determinant of adverse cardiac remodelling and whole heart contractility. Expression of the transcription factor RUNX1 is increased in the border zone at 1 day after myocardial infarction, suggesting potential for targeted therapeutic intervention. Here we demonstrate that RUNX1 drives reductions in cardiomyocyte contractility, sarcoplasmic reticulum-mediated calcium release, mitochondrial density, and the expression of genes important for oxidative phosphorylation.

Antagonising RUNX1 expression *via* short-hairpin RNA interference preserved cardiac contractile function following myocardial infarction when delivered either via direct adenoviral delivery into the border zone or *via* an adeno-associated virus vector administered intravenously. Equivalent effects were obtained with a small molecule inhibitor (Ro5-3335) that reduces RUNX1 function by blocking its interaction with the essential co-factor CBFβ. Both tamoxifen-inducible *Runx1*-deficient and *Cbfβ*-deficient cardiomyocyte-specific mouse models demonstrated that antagonising RUNX1 function preserves the expression of genes important for oxidative phosphorylation following myocardial infarction. Our results confirm the translational potential of RUNX1 as a novel therapeutic target in myocardial infarction, with wider opportunities for use across a range of cardiac diseases where RUNX1 drives adverse cardiac remodelling.

## INTRODUCTION

Myocardial infarction (MI) due to acute coronary artery blockage leads to cardiomyocyte death and an injury response culminating in the generation of three regions within the left ventricle (LV). The infarct zone (IZ) predominantly comprises fibrillar collagens that maintain the integrity of the myocardium and prevent LV wall rupture. The remote zone (RZ), is located furthest away from the IZ but over time develops several cellular and extracellular matrix changes that impact LV function. The border zone (BZ) surrounds the IZ; myocardium in this region is viable and perfused but hypocontractile. Cellular changes within these three regions are fundamental to the process of adverse cardiac remodelling, which manifests clinically as LV wall thinning, dilation and reduced contractility as early as 1 day post-MI^1–6^. Together with neurohumoral activation, adverse cardiac remodelling post-MI can lead to the clinical syndrome of heart failure (HF) with reduced ejection fraction, which despite optimised medical and device therapy is associated with high mortality rates^7^.

Cellular changes that occur in the BZ during the first few days post-MI play a critical role in the progression of LV remodelling, contractility, and patient prognosis during the subsequent weeks and months^6, 8–10, 11, 12^. Many of these BZ changes are conserved between mice and humans and include abnormal sarcoplasmic reticulum (SR)-mediated calcium release^1, 2, 13–16^ and the downregulation of genes important for mitochondrial oxidative phosphorylation^17^. The identification of drivers that induce these early changes in the BZ are critical for the development of new therapeutic approaches to prevent progression of adverse cardiac remodelling and development of HF following MI.

*RUNX1* encodes a DNA-binding α-subunit that partners with a common β subunit (CBFβ) to form a heterodimeric transcription factor that acts as both an activator and repressor of target genes in normal development and disease^18^. Although RUNX1 has been intensively studied in cancer and haematology, its role in the heart is only just emerging^19^. *Runx1* expression is increased in cardiomyocytes located within the BZ and IZ region as early as 1 day post-MI and mediates impaired contractility^20–22^. Whether this increased RUNX1 expression in the BZ can be therapeutically targeted to preserve contractility following MI remains unknown.

In this study, we address this knowledge gap by: (i) determining that the spatially restricted increased expression of RUNX1 in the BZ drives impaired calcium handling, a decrease in mitochondrial density, and reduced expression of genes important for oxidative phosphorylation in the BZ; and (ii) revealing that multiple therapeutic approaches can be used to inhibit Runx1 in the BZ and thereby protect myocardial contractility following MI.

## RESULTS

### Calcium handling in BZ and RZ cardiomyocytes at 1 day post-MI

SR-mediated calcium release leads to sarcomere shortening and contraction, which are key processes determining LV pump function. Differences in BZ and RZ cardiomyocyte SR-mediated calcium release in the first week post-MI contribute to regional heterogeneity of contractile function across the LV. This regional heterogeneity contributes to impaired global LV contractile function and poor patient prognosis^8–10^. *Runx1* expression in BZ cardiomyocytes is increased as early as day 1 following MI^20^. Whether this change in *Runx1* expression at this time point simply coincides with, or impacts on, SR-mediated calcium handling was unknown. Therefore, we compared SR calcium handling in BZ and RZ cardiomyocytes isolated from the hearts of C57BL/6J and tamoxifen-inducible cardiomyocyte-specific *Runx1*-deficient (*Runx1*^Δ/Δ^) mice (and in Cre negative *Runx1*^fl/fl^ littermate controls)^20^ at 1-day post-MI (Fig.1a). These results were compared to cardiomyocytes isolated from C57BL/6J hearts before MI at equivalent regions to where the “BZ” and “RZ” would be expected post-MI. Cardiomyocytes were stimulated at 1.0 Hz to elicit SR-mediated calcium release into the cytosol (calcium transients), the amplitude of which largely determines the force of contraction (Fig.1a).

**Figure 1.**
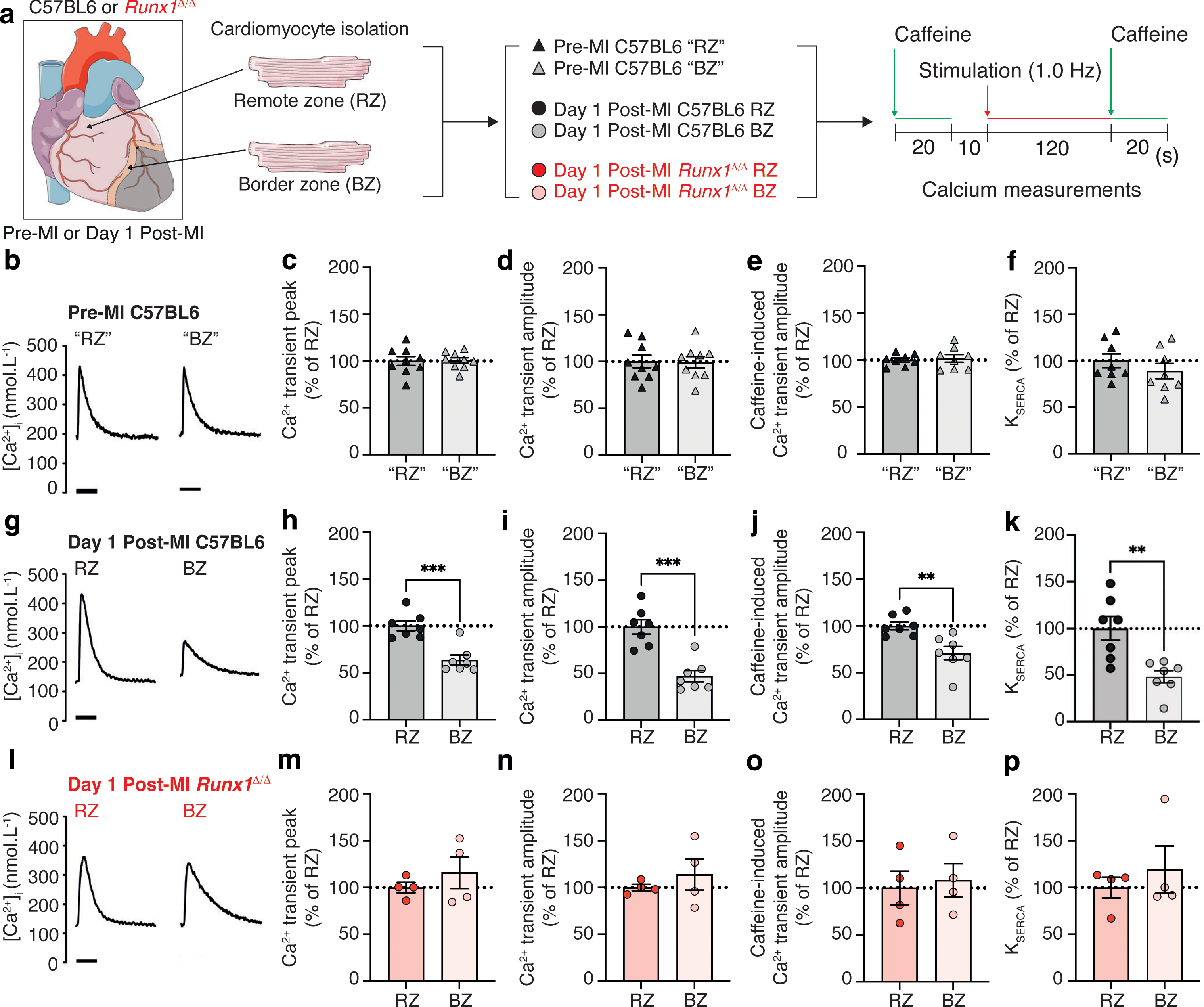
Excitation-contraction coupling in C57BL/6J and *Runx1*^Δ/Δ^ mice and 1 day post-myocardial infarction (MI). **(a)** Protocol. **(b)** Typical calcium (Ca^2+^) transients before MI in C57BL/6J mice in the “remote zone” (“RZ”) and “border zone” (“BZ”). **(c)** Mean Ca^2+^ transient peak, **(d)** mean Ca^2+^ transient amplitude *n*=76 (9 hearts) and from the BZ *n*=43 (9 hearts), **(e)** Mean caffeine-induced Ca^2+^ transient amplitude and **(f)** mean sarcoendoplasmic reticulum calcium transport ATPase (SERCA) activity from C57BL/6J mice from RZ, *n*=76 (8 hearts) and from the BZ *n*=43 (8 hearts). **(g)** Typical Ca^2+^ transients 1 day post-MI in C57BL/6J mice in the RZ and BZ. **(h)** Mean Ca^2+^ transient peak, **(i)** mean Ca^2+^ transient amplitude, **(j)** mean caffeine-induced Ca^2+^ transient amplitude and **(k)** mean SERCA activity from C57BL/6J 1-day post-MI mice from RZ, *n*=64 (7 hearts) and from the BZ *n*=30 (7 hearts). **(l)** Typical Ca^2+^ transients 1-day post-MI in *Runx1^Δ/Δ^* mice in the RZ and BZ. **(m)** Mean Ca^2+^ transient peak, **(n)** mean Ca^2+^ transient amplitude, **(o)** mean caffeine induced Ca^2+^ transient amplitude and **(p)** mean SERCA activity from *Runx1^Δ/Δ^* mice 1-day post-MI mice from RZ, *n*=16 (4 hearts) and from the BZ *n*=9 (4 hearts). Error bars represent mean ± SEM. \*\**P*<0.01, \*\*\**P*<0.001.

By contrast to hearts without MI (Fig.1b and 1c), the calcium transient peak (systolic [Ca^2+^]i) following MI in BZ cardiomyocytes was 64% of the peak in RZ cardiomyocytes (*P*<0.05; Extended Table 1a-c; Fig. 1g and 1h). In *Runx1*^Δ/Δ^ hearts, the calcium transient peak was preserved in BZ cardiomyocytes relative to RZ cardiomyocytes (*P*>0.05; Fig. 1l and 1m). There was no difference in the diastolic [Ca^2+^]i (calcium transient minimum) between BZ and RZ in all groups (*P*>0.05; Extended Fig.1). Unlike hearts without MI (Fig. 1b and 1d), the changes in systolic [Ca^2+^]i in BZ cardiomyocytes following MI resulted in a calcium transient amplitude which was 47% of that observed in RZ cardiomyocytes (*P*<0.05; Fig.1g and 1i). By contrast, the calcium transient amplitude in BZ cardiomyocytes of *Runx1^Δ/Δ^* hearts was preserved (*P*>0.05; Fig. 1l and 1n; Extended Table 1).

The SR calcium content is the predominant determinant of calcium transit amplitude^23, 24^. To determine the SR calcium content, a rapid bolus of caffeine (10 mmol/L) was applied at the end of the protocol (Fig.1a) to release all the calcium from the SR into the cytosol, permitting the quantification of the SR calcium content. In contrast to hearts without MI (Fig.1e), the caffeine-induced Ca^2+^ transient amplitude (SR calcium content) of BZ cardiomyocytes at 1 day post-MI was 71% of that observed in RZ cardiomyocytes (*P*<0.05; Fig. 1j; Extended Table 1). The SR calcium content of *Runx1*^Δ/Δ^ BZ cardiomyocytes 1 day post-MI was not significantly different to that from RZ cardiomyocytes of the same hearts (*P*>0.05; Fig. 1o).

A key determinant of the SR calcium content is uptake into the SR via the calcium ATPase pump SERCA^25^. We hypothesised that the lowered SR calcium content in BZ cardiomyocytes following MI might reflect decreased SERCA activity (K_SERCA_) or enhanced extrusion of calcium from the cell via the sodium-calcium exchanger. To determine K_SERCA_, we measured the rate constant of decay of the caffeine-induced calcium transient (which includes sarcolemmal efflux but not SR calcium uptake) and subtracted this value from that of the electrically stimulated calcium transient (which includes both SR calcium uptake and sarcolemmal efflux)^26, 27^. In contrast to hearts without MI (Fig.1f), K_SERCA_ of BZ cardiomyocytes following MI was 48% of RZ cardiomyocytes (*P*<0.05; Fig. 1k; Extended Table 1). However, SERCA activity was not different between *Runx1*^Δ/Δ^ BZ and RZ cardiomyocytes 1 day following MI (*P*>0.05; Fig.1p). Extrusion of calcium from the cell via the sodium-calcium exchanger was assessed by the time constant of caffeine-induced calcium transient decay but was not different between the BZ and RZ cardiomyocytes of any group (Extended Table 1).

Separate experiments confirmed that the relative difference between BZ and RZ in all parameters measured in C57BL6J mice at 1-day post-MI (Fig.1g-k) were also observed in *Runx1^fl^*^/fl^ mice at 1 day post-MI (i.e. the Cre-negative littermate controls for *Runx1*^Δ/Δ^ mice; Extended Table 1).

These data demonstrate that marked differences in SR-mediated calcium release/uptake between BZ and RZ cardiomyocytes that are known to contribute to whole heart contractile dysfunction in the first week post-MI, are evident as early as 1-day post-MI in both the C57BL/6J and littermate controls and are *Runx1*-dependent.

### RNA-sequencing analyses of BZ and RZ myocardium in *Runx1*^Δ/Δ^ mice

The BZ is a discrete and highly active region of the myocardium post-MI that undergoes a considerable number of transcriptional changes, some of which ultimately may drive adverse cardiac remodelling. The extent to which the observed increase of *Runx1* within the BZ plays a role in these transcriptional changes was unknown. Therefore, we used RNAseq to compare the changes in gene expression in the BZ myocardium relative to the RZ in both *Runx1*^Δ/Δ^ and *Runx1*^fl/fl^ control mice at 1 day following MI. Using a false discovery rate (FDR) cut-off of ≤ 0.05, there were 7166 differentially expressed genes in the BZ relative to the RZ in *Runx1*^fl/fl^ control mice (Fig. 2a). By contrast, there were only 1748 differentially expressed genes in the BZ relative to the RZ in *Runx1*^Δ/Δ^ mice. There were 1618 differentially expressed genes that were common to both *Runx1*^Δ/Δ^ and *Runx1^fl^*^/fl^ control mice leaving 130 differentially expressed genes unique to *Runx1*^Δ/Δ^ mice (Fig. 2a and 2b).

**Figure 2.**
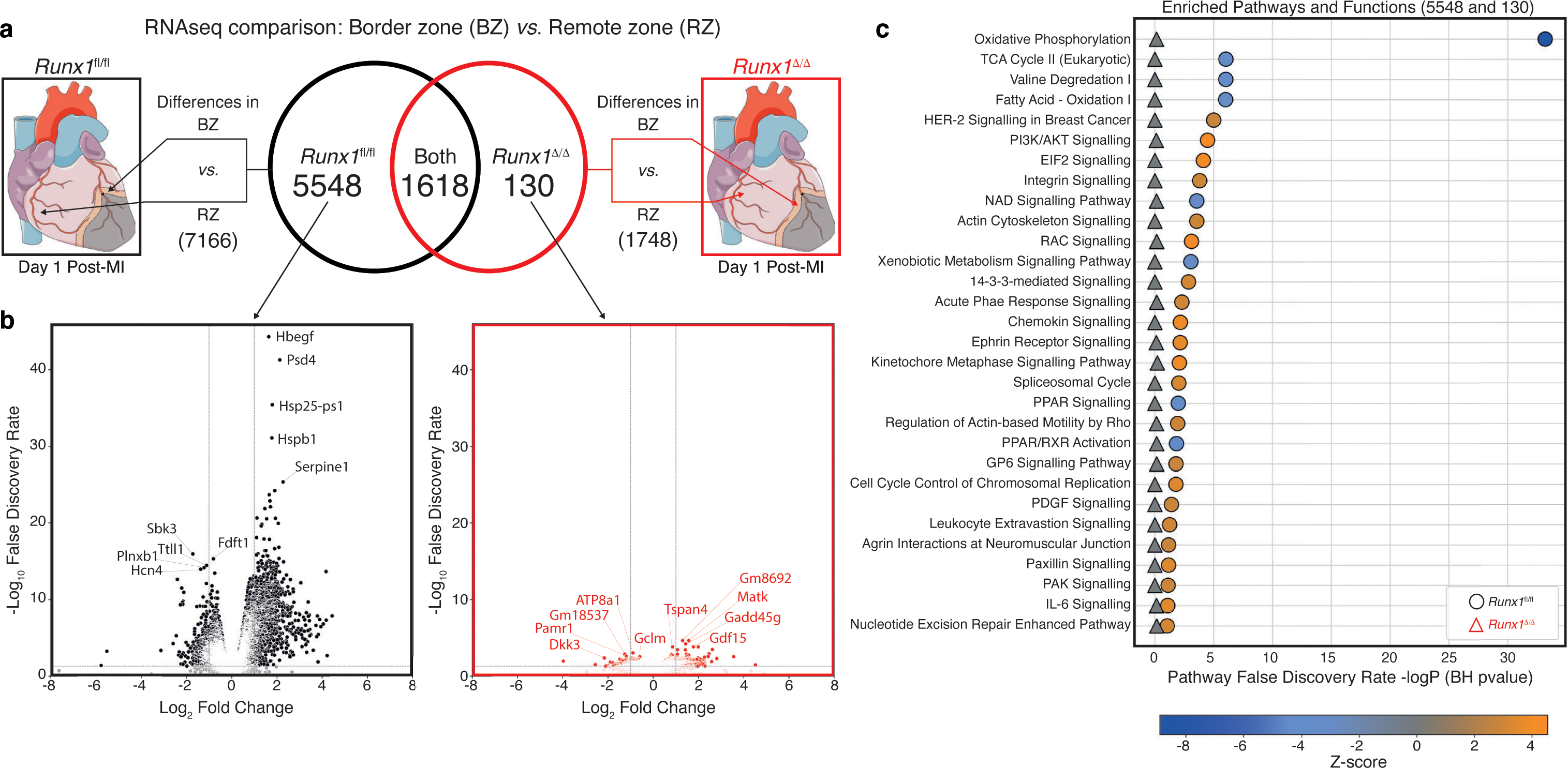
RNA sequencing comparisons and analysis between border zone (BZ) and remote zone (RZ) in *Runx1*^fl/fl^ and *Runx1*^Δ/Δ^ mice 1 day post-myocardial infarction (MI). **(a)** Schematic of comparison and Venn diagram of gene differences. **(b)** Volcano plots of differentially regulated genes unique to *Runx1*^fl/fl^ mice (left, black) and *Runx1*^Δ/Δ^ mice (right, red) in the BZ versus the RZ at 1 day post-MI. Top five upregulated and down regulated genes based on false discovery rate (FDR) are noted. **(c)** Enriched biological pathways ranked by logP (B-H p value) using IPA analysis from unique differences in *Runx1*^fl/fl^ mice (circles) compared with BZ and RZ differences in *Runx1*^Δ/Δ^ mice (triangles). Blue-to-orange heatmap and symbol colours represent predicted inhibition (blue) and activation (orange) or no change (grey) of pathways based on Z-score.

Ingenuity pathway analysis (IPA) software was used to determine the enriched pathways and functions specific to the 5548 differentially expressed genes unique to *Runx1*^fl/fl^ mice, as compared to the 130 differentially expressed genes unique to the *Runx1*^Δ/Δ^ mice. Oxidative phosphorylation was the most highly significant changed pathway/function (Fig. 2c).

Further interrogation of these pathways using IPA analysis of the total number of differentially expressed genes (between the BZ and RZ) in *Runx1*^fl/fl^ and *Runx1*^Δ/Δ^ post-MI, revealed marked downregulation of genes involved in oxidative phosphorylation across all five inner mitochondrial complexes in BZ myocardium of *Runx1*^fl/fl^ control mice (Fig. 3a and Extended Fig. 2). By contrast, no genes involved in oxidative phosphorylation within mitochondria were significantly downregulated in the BZ of *Runx1*^Δ/Δ^ mice suggesting that *Runx1* deficiency within cardiomyocytes preserves genes involved in oxidative phosphorylation within the BZ following MI (Fig. 3b).

**Figure 3.**
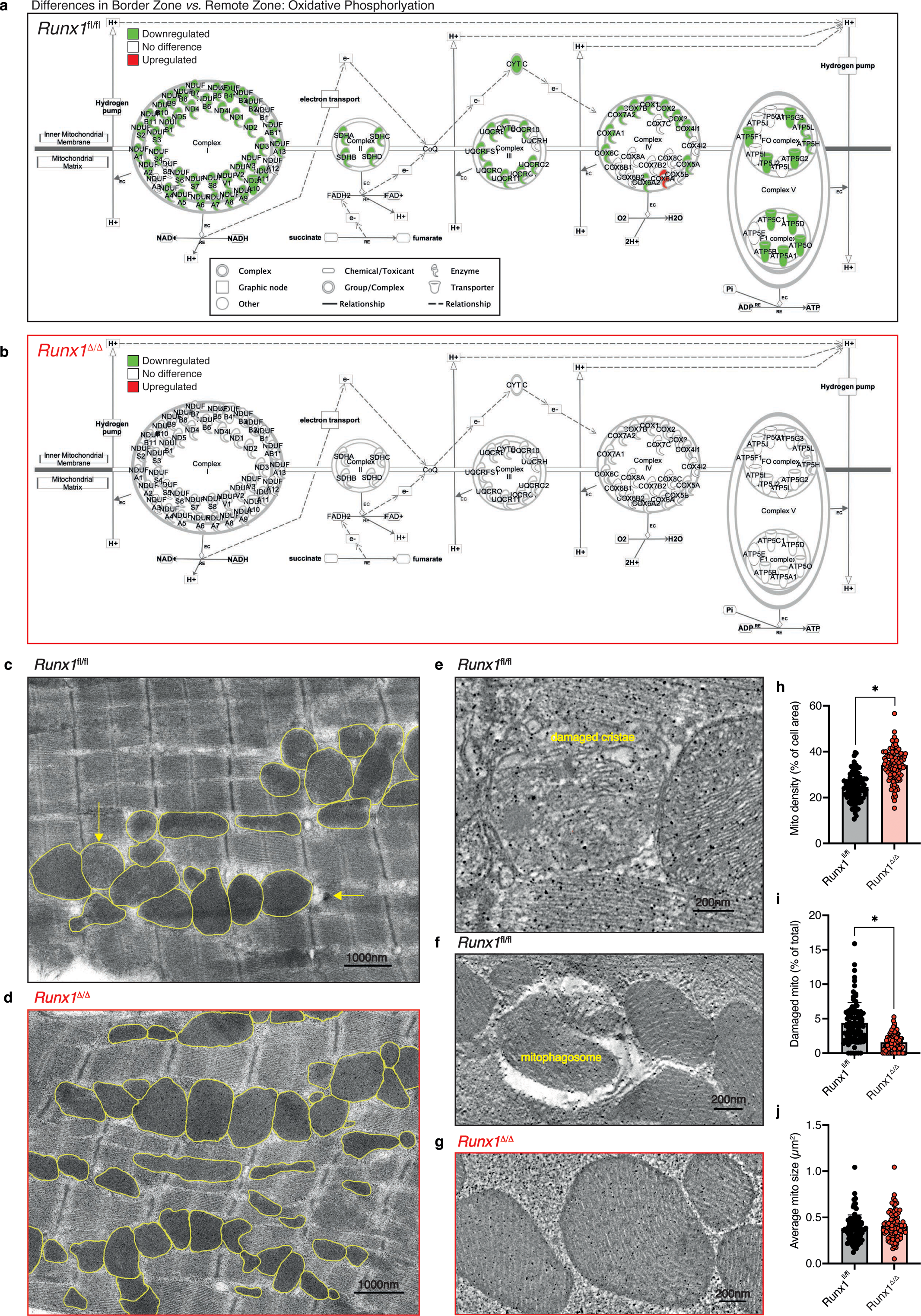
Oxidative phosphorylation and mitochondrial density in the border zone (BZ) of Runx1^fl/fl^ and *Runx1*^Δ/Δ^ mice 1 day post-myocardial infarction (MI). Schematic visualisation of genes involved in oxidative phosphorylation complexes from the RNAseq data from the border zone (BZ) and remote zone (RZ) in *Runx1*^fl/fl^ and *Runx1*^Δ/Δ^ mice 1 day post-MI in Ingenuity Pathway Analysis (IPA) (downregulated – green, upregulated – red, no difference – white) of **(a)** *Runx1*^fl/fl^ mice (7166 differentially expression genes) and **(b)** *Runx1*^Δ/Δ^ mice (1748 differentially expressed genes). Low resolution electron microscopy images from BZ of **(c)** *Runx1*^fl/fl^ mice and **(d)** *Runx1*^Δ/Δ^ mice. Mitochondria are outlined in yellow, with damaged mitochondria indicated by yellow arrows. High-resolution electron tomography representative images of **(e-f)** damaged mitochondria from the BZ of *Runx1*^fl/fl^ mice 1 day post-MI and **(g)** healthy mitochondria from the BZ of *Runx1*^Δ/Δ^ mice 1 day post-MI. Quantification of **(h)** mitochondrial density as a percentage of cell area, **(i)** the number of damaged mitochondria as a percentage of the total number of mitochondria, and **(j)** average mitochondrial size from electron microscopy images of the BZ from *Runx1*^fl/fl^ mice and *Runx1*^Δ/Δ^ mice 1 day post-MI *n*=92 (2 hearts). Error bars represent mean ± SEM. \**P*<0.05, unpaired Student t-test on average heart data.

Mitochondria are critical for the energy production of the heart in the form of ATP generation via the process of oxidative phosphorylation. Given the marked preservation of expression of genes involved in oxidative phosphorylation within the BZ of *Runx1*^Δ/Δ^ mice, we next quantified mitochondrial density, integrity, and size from electron microscopy images taken from the BZ region of *Runx1*^fl/fl^ and *Runx1*^Δ/Δ^ mice at 1-day post-MI. The BZ of *Runx1*^fl/fl^ mice had reduced mitochondrial density compared to the BZ of *Runx1*^Δ/Δ^ mice (24% and 34% of cell area respectively, *P*<0.05; Fig. 3c, 3d and 3h). There were significantly more damaged mitochondria (identified by dissolved/damaged cristae or mitophagosomes) in the BZ of *Runx1*^fl/fl^ mice (4.4% of all mitochondria) compared to the BZ of *Runx1*^Δ/Δ^ mice (1.5% of all mitochondria, *P*<0.05; Fig. 3e-g and 3i). No difference in mitochondrial size was detectable between groups (Fig. 3j).

### Targeted BZ knockdown of *Runx1* by injection of adenoviral-*Runx1*-shRNA post-MI

We have previously shown that genetically modified cardiomyocyte-specific *Runx1*-deficient mice (*Runx1*^Δ/Δ^) have preserved left ventricle (LV) cardiac contractility following MI^20^. However, whether increased *Runx1* expression post-MI in the BZ cardiomyocytes can be therapeutically targeted remained unknown. To address this gap in our knowledge we utilised various approaches. The first approach was to inject an adenoviral (Ad) vector expressing *Runx1*-shRNA (or control Ad-scrambled-shRNA) directly into the BZ area of C57BL/6J mice immediately following MI (Fig. 4a) to reduce *Runx1* expression and determine the impact on LV contractility. The ability of the adenovirus to reduce *Runx1* expression was first verified in IP1B cells, a murine cell line with high *Runx1* expression (Extended Fig. 3). We next determined the efficiency of transduction and *Runx1* expression (RNA hybridisation using RNAscope, as previously published^20^) in cardiomyocytes (identified by PCM-1 expression) and other cardiac cells (Fig. 4b). There was no change in the total number of cardiomyocyte nuclei or non-cardiomyocyte nuclei between control and Ad-*Runx1*-shRNA groups within the IZ, BZ and RZ (Extended Fig. 4). Hearts injected with the control Ad-scrambled-shRNA demonstrated the expected increase in *Runx1* expression within the IZ and BZ region of the LV; whereas *Runx1* expression within the RZ of the LV remained low (Fig. 4c). Injection of Ad-*Runx1*-shRNA into the BZ resulted in a 46% reduction in *Runx1* expression within cardiomyocyte nuclei relative to Ad-scramble-shRNA injected hearts (39.1±4.5 *vs*. 72.3±3.1% of total number of cardiomyocytes, *P*<0.05; Fig. 4c). The reduction in *Runx1* expression in cardiomyocytes was specific to the BZ region as there was no change in *Runx1* expression within cardiomyocyte nuclei between the two groups in the other LV regions (Fig. 4c).

**Figure 4.**
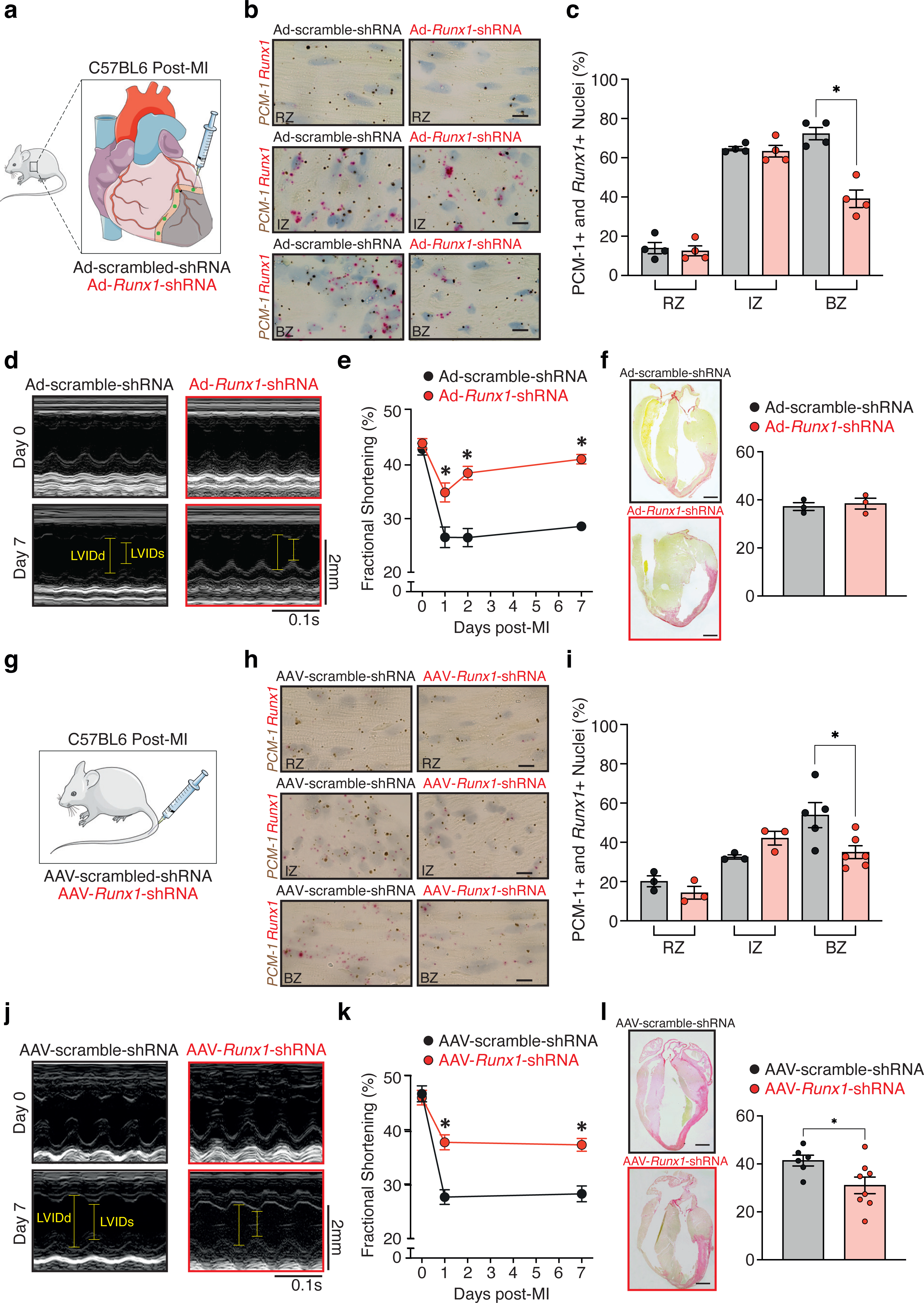
Cardiac function and *Runx1* expression in Ad-*Runx1*-shRNA and AAV-*Runx1-*shRNA mice. (**a**) Schematic representation of Ad given *via* an injection into the border zone immediately after induction of MI. (**b-e**) Typical images of regional heart sections by RNA i*n situ* hybridisation (using RNAscope). Regions examined were the remote zone (RZ), infarct zone (IZ) and border zone (BZ) at 7 days post-MI in Ad-*Runx1*-shRNA injected mice (*n*=4) and Ad-scramble-shRNA injected mice (*n*=4). Probes for *Runx1* (pink punctate dots) and pericentriolar material 1 (*PCM-1*) (brown punctate dots) were used. Scale bar, 10 µm; magnification 100x. (**c**) Mean quantification of *Runx1-*positive cardiomyocyte nuclei (*PCM-1*+ and *Runx1*) expressed as a percentage of total nuclei (data from b). \**P*<0.05, 7 days after MI Ad-*Runx1*-shRNA *vs.* Ad-scramble-shRNA. (**d**) Echocardiography (scale: x=0.1 s; y=2 mm). (**e**) The 7-day echocardiographic data for fractional shortening (FS) of Ad-*Runx1*-shRNA (*n*=8; day 0, *n*=8; day 1, *n*=7; day 2, *n*=7; day 7) *vs.* Ad-scramble-shRNA (n=8; day 0, *n*=8; day 1, *n*=7; day 2, *n*=7; day 7). (**f**) Typical picrosirius-red-stained hearts and infarct size as the percentage of the left ventricle (*n*=3). (**g**) Schematic representation of AAV given *via* tail vein injection (**h**) Typical images of regional heart sections by RNA i*n situ* hybridisation (using RNAscope). Regions examined were the RZ (AAV-*Runx1*-shRNA [*n*=3] vs. AAV-scramble-shRNA injected mice [*n*=3]), IZ (AAV-*Runx1*-shRNA [*n*=3] vs. AAV-scramble-shRNA injected mice [*n*=3]), and BZ (AAV-*Runx1*-shRNA [*n*=6] vs. AAV-scramble-shRNA injected mice [*n*=5]), at 7 days post-MI. Probes for *Runx1* (pink punctate dots) and pericentriolar material 1 (*PCM-1*) (brown punctate dots) were used. Scale bar, 10 µm; magnification 100x. (**i**) Mean quantification of *Runx1* positive cardiomyocyte nuclei (*PCM-1*+ and *Runx1*) expressed as a percentage of total nuclei (data from h). *= *P*<0.05, 7 days after MI AAV-*Runx1*-shRNA *vs.* AAV-scramble-shRNA. (**j**) Echocardiography (scale: x=0.1 s; y=2 mm). (**k**) The 7-day echocardiographic data for fractional shortening (FS) (AAV-*Runx1*-shRNA [*n*=9; day 0, *n*=9; day 1, *n*=9; day 7] *vs.* AAV-scramble-shRNA [*n*=10; day 0, *n*=10; day 1, *n*=9; day 7]). (**l**) Typical picrosirius-red stained hearts and infarct size as the percentage of the left ventricle AAV-*Runx1*-shRNA injected mice (*n*=8) *vs.* AAV-scramble-shRNA injected mice (*n*=6)). Error bars represent mean ± SEM. * *P*<0.05, Student *t* test.

Echocardiography was performed before MI and at 1 , 2 and 7 days post-MI to assess LV function (Fig. 4d). As expected, at 1 day post-MI there was a decline in cardiac systolic function (measured by fractional shortening; %FS) in the Ad-scramble-shRNA injected control group of mice which was maintained at a low level from day 1-7. By contrast, the Ad-*Runx1*-shRNA group had a markedly preserved contractile function at all time points post-MI relative to the Ad-scramble-shRNA injected control group (day 1, 34.7±1.8 *vs*. 26.4 ±1.9; day 2, 38.3±1.3 *vs*.; 26.4±1.7 and day 7, 40.9±0.9 *vs*. 28.5±0.6% FS; *P*<0.05; Fig. 4e). The preservation of fractional shortening in C57BL/6J post-MI mice treated with Ad-*Runx1*-shRNA was attributable to an improved contractility as evidenced by a smaller LV internal diameter (LVIDs) measured during systole at all time points post-MI relative to the Ad-scramble-shRNA injected control group (day 1, 2.4±0.1 *vs*. 3.0±0.1; day 2, 2.3±0.1 *vs*. 3.1±0.1 *vs*. day 7; 2.3±0.2 *vs*. 2.8±0.1mm LVIDs; *P*<0.05; Extended Table 2). The hearts of C57BL/6J mice treated with Ad-*Runx1*-shRNA were less dilated at 1 and 2 days post-MI as measured by the LV internal diameter measured during diastole (LVIDd) but was not different at 7 days post-MI between the groups (day 1, 3.7±0.1 *vs*. 4.1±0.2 [*P*<0.05]; day 2, 3.7 ±0.2 *vs*. 4.3±0.1 [*P*<0.05] day 7; 3.9±0.2 *vs*. 3.9±0.2mm [*P*>0.05]; Extended Table 2).

To determine whether a change in infarct size contributed to preserved cardiac function in C57BL/6J mice treated with Ad-*Runx1*-shRNA following MI, Sirius red staining was performed on heart slices (Fig. 4f). Infarct size was not different between the Ad-*Runx1*-shRNA and Ad-scramble-shRNA injected control group (*P*>0.05; Fig. 4f).

### Cardiac function and structure, and localised expression of *Runx1* following treatment with AAV9-*Runx1*-shRNA post-MI

Although knockdown of *Runx1* in the BZ myocardium was effective in preserving cardiac contractile function, the need to directly inject the heart with the Ad may have reduced the translational potential of the approach. We, therefore, explored the use of a cardiotropic adeno-associated virus serotype 9 (AAV9) expressing a short-hairpin RNA targeting *Runx1,* which can be injected intravascularly to knockdown *Runx1* within the heart. AAV vectors are widely used for cardiac gene delivery in pre-clinical models and clinical trials^28^ and a licensed AAV9 gene therapy is available for treatment of spinal muscular atrophy^29^. AAV9 encoding shRNA provides highly efficient knockdown in the heart^30^. AAV9-*Runx1*-shRNA or control AAV-scramble-shRNA were delivered via tail-vein injection following MI in C57BL/6J mice (Fig. 4g). Using RNAscope, we determined the expression of *Runx1* within AAV9-*Runx1*-shRNA injected MI hearts relative to AAV9-scramble-shRNA within RZ, BZ and IZ of each heart at day 7 post-MI (Fig. 4h). *Runx1* expression within cardiomyocyte nuclei of the BZ of mice injected with AAV9-*Runx1*-shRNA was 65% of the levels observed in mice injected with AAV9-scramble-shRNA (35.0±3.6 *vs*. 53.9±6.4% of total number of cardiomyocytes; P<0.05; Fig. 4i). Despite a decrease in mean levels of *Runx1* expression within cardiomyocyte nuclei in the RZ of mice injected with AAV9-*Runx1*-shRNA relative to mice injected with AAV9-scramble-shRNA, the observed change was not significant (14.3±2.3 *vs*. 20.1±2.8; *P*>0.05; Fig. 4i). No significant change in *Runx1* expression in cardiomyocytes was found in the IZ region (Fig. 4i). There was no change in the total number of cardiomyocyte nuclei or non-cardiomyocyte nuclei between control and AAV-*Runx1*-shRNA groups within the three regions assessed (Extended Fig. 4).

Echocardiography was performed before MI (day 0) and at 1 day and 7 days following MI to assess LV function (Fig. 4j). As expected, at 1 day post-MI there was a decline in cardiac systolic function (measured by fractional shortening; %FS) in the AAV9-scramble-shRNA injected control group of mice which was maintained at a low level from days 1-7. By contrast, the AAV9-*Runx1*-shRNA group had a markedly preserved contractile function at day 1 and day 7 post-MI relative to the AAV9-scramble-shRNA injected control group (day-1, 37.7±1.4 *vs*. 27.6±1.4; day 7; 37.2±1.2 *vs*. 28.2±1.5% FS; *P*<0.05, Fig. 4k). The preservation of fractional shortening in C57BL/6J-MI mice treated with AAV9-*Runx1*-shRNA was attributable to an improved contraction as evidenced by a smaller LVIDs measured during systole at day 1 and day 7 post-MI relative to the AAV9-scramble-shRNA injected control group (day-1, 2.4±0.1 *vs*. 2.9 ±0.2; day 7, 2.5±0.1 *vs*. 3.1±0.2mm; *P*<0.05; Extended Table 2). Cardiac dilation was not significantly affected by AAV9-*Runx1*-shRNA as LV internal diameter measured during diastole was not different at day 1 and day 7 post-MI (Extended Table 2).

In contrast to direct injection of Ad*Runx1*-shRNA into the BZ, intravenous injection of *AAV9-Runx1*-shRNA reduced infarct size in C57BL/6J-MI mice relative to AAV9-scramble-shRNA group post-MI (41.4±1.9 *vs*. 31.1±3.5% of LV; *P*<0.05; Fig. 4l).

### Small molecule mediated inhibition of *Runx1* post-MI

RUNX1 partners with CBFβ to act as a heterodimeric transcription factor^18^. Interfering with RUNX1 binding to CBFβ significantly reduces affinity for RUNX1 binding to its target genes thus its transcriptional activity. The benzodiazepine derivative Ro5-3335 is an established RUNX1 inhibitor that alters the interaction between RUNX1 and CBFβ^31^. We used Ro5-3335 in two different protocols as an alternative therapeutic approach to gene transfer in the context of MI (Fig. 5a). The first protocol involved pre-treatment with Ro5-335 in C57BL/6J mice every other day for 7 days followed by the same treatment pattern following MI. Echocardiography was performed before MI (day 0) and at 1, 3, and 7 days following MI to assess LV function (Fig. 5b). At 1 day post-MI there was a decline in cardiac systolic function (measured by %FS) in the vehicle-injected group of mice that was maintained at a low level from day 1 to 7. By contrast, the Ro5-3335 injected group had a preserved contractile function at days 1 and 7 points post-MI relative to the vehicle control group (day 1, 37.6±2.2 *vs*. 30.1±0.6; day 3, 40.4±3.5 *vs*. 32.8±1.2; day 7; 38.8±2.0 *vs*. 30.1±2.2% FS; *P*<0.05, Fig. 5b).

**Figure 5.**
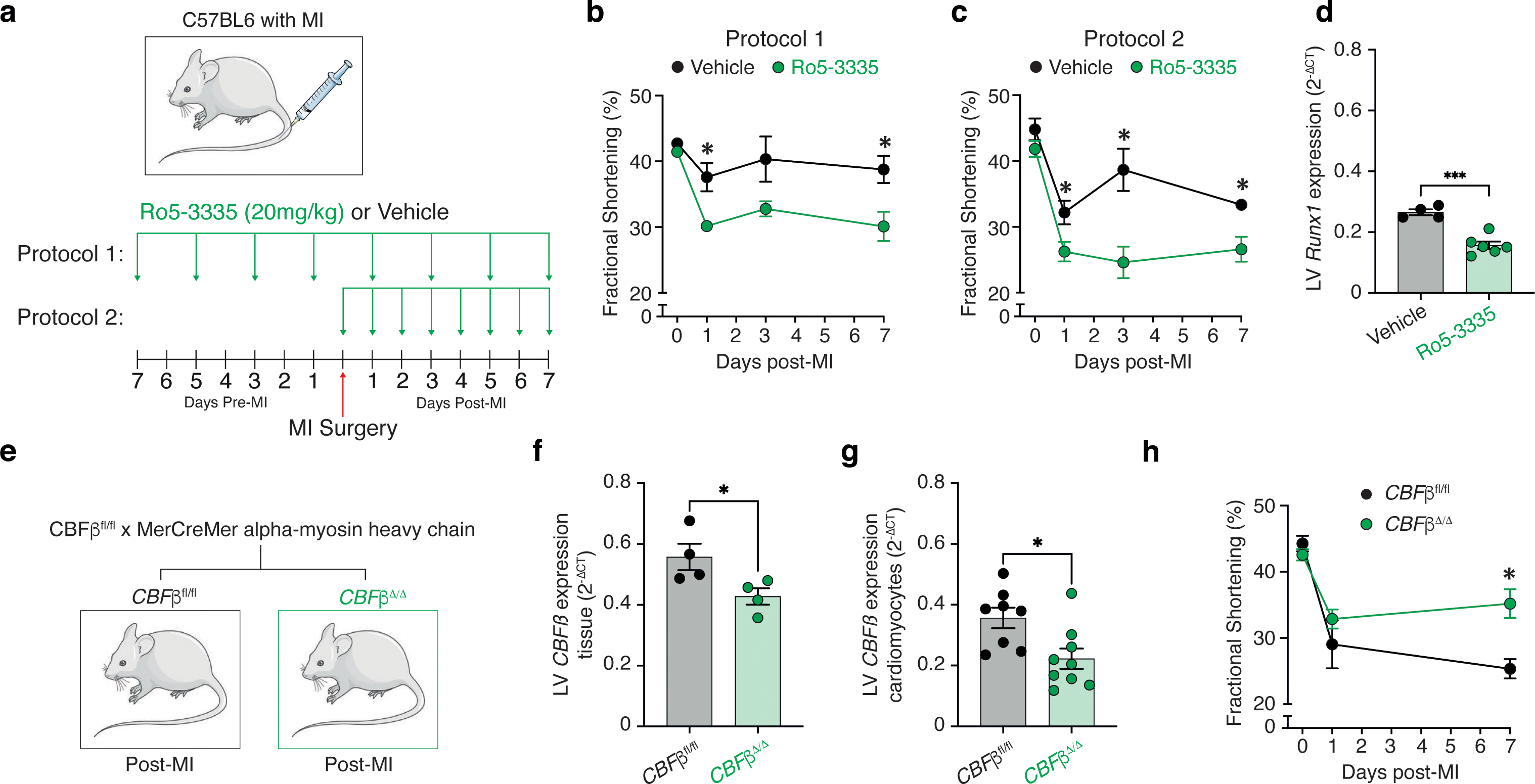
Cardiac function and *Runx1* expression in Ro5-3335 treated MI mice and *Cbfβ* ^Δ/Δ^ MI mice. **(a)** Protocol. (**b**) The 7-day echocardiographic data for fractional shortening (FS) for mice receiving protocol 1: Ro5-3335-treated MI mice (*n*=6; day 0, *n*=6; day 1, *n*=5; day 3, *n*=5; day 7) *vs.* vehicle (DMSO)-treated MI mice (*n*=6; day 0, *n*=5; day 1, *n*=3; day 3, *n*=4; day 7). (**c**) The 7-day echocardiographic data for FS for mice receiving protocol 2: Ro5-3335-treated mice (*n*=8; day 0, *n*=8; day 1, *n*=6; day 3, *n*=6; day 7) *vs.* vehicle (DMSO)-treated mice (*n*=7; day 0, *n*=8; day 1, *n*=3; day 3, *n*=7; day 7). \**P*<0.05, Student *t* test. (**d**) *Runx1* mRNA expression relative to Peptidylprolyl Isomerase B (*Ppib)* as measured by real-time quantitative polymerase chain reaction (qPCR) in LV tissue from Ro5-3335-treated MI mice (*n*=6) *vs.* vehicle (DMSO)-treated MI mice (*n*=4). \*\*\**P*<0.005, Student *t* test. (**e**) Protocol. (**f**) *Runx1* mRNA expression relative to *Ppib* as measured by qPCR in LV tissue from *Cbfβ* ^Δ/Δ^ MI mice (*n*=4) *vs. Cbfβ^fl/fl^* MI mice (*n*=4). (**g)** *Runx1* mRNA expression relative to *Ppib* as measured by qPCR in LV isolated cardiomyocytes from *Cbfβ* ^Δ/Δ^ MI mice (*n*=9) *vs. Cbfβ^fl/fl^* MI mice (*n*=8). \**P*<0.05, Student *t* test. (**h**) The 7-day echocardiographic data for FS for from *Cbfβ* ^Δ/Δ^ MI mice (*n*=8; day 0, *n*=6; day 1, *n*=6; day 7) *vs. Cbfβ^fl/fl^* MI mice (*n*=6; day 0, *n*=;5 day 1, *n*=6; day 7). Error bars represent mean ± SEM. \**P*<0.05, Student *t* test.

The second protocol involved post-treatment with Ro5-3335 in C57BL/6J mice every day for 7 days following MI. Echocardiography was performed before MI (day 0) and at 1, 3, and 7 days following MI to assess LV function (Fig. 5a; Extended Table 2). At 1 day post-MI there was a decline in cardiac systolic function (measured by %FS) in the vehicle injected group of mice that was maintained at a low level from day 1 to day 7 (Fig. 5c). By contrast, the Ro5-3335 injected group had a markedly preserved contractile function at all time points post-MI relative to the vehicle control group (day 1, 32.2±1.8 *vs*. 26.2±1.5; day 3, 38.6±3.2 *vs*. 24.6±2.4; day 7; 33.3±0.6 *vs*. 26.6±1.9% FS; *P*<0.05, Fig. 5c).

As observed in previous studies that have used Ro5-3335 to inhibit Runx1 in other tissues^32^, *Runx1* expression in LV myocardium was decreased relative to vehicle control (*P*<0.05; Fig. 5d).

These data confirm that both gene therapeutic and pharmacological approaches to inhibit Runx1 preserve cardiac contractile function following MI.

### Generation of a cardiomyocyte-specific *Cbfβ* deficient mouse to inhibit RUNX1 function

We next sought to confirm our results with Ro5-3335 using an organ-specific approach to limit the potential for CBFβ to interact with RUNX1. This was achieved by generating a tamoxifen-inducible cardiomyocyte-specific CBFβ deficient mouse using the Cre-LoxP system. *Cbfβ*^fl/fl^ mice, described previously^33^, were crossed with mice expressing tamoxifen-inducible Cre recombinase (*MerCreMer*) under the control of the cardiac-specific *αMHC* (α-myosin heavy chain)^34^ to produce the relevant test (*Cbfβ*^Δ/Δ^) and littermate control (*Cbfβ^fl/fl^*) cohorts (Fig. 5e). Expression of *Cbfβ* was reduced to 77% and 47% in LV myocardium and isolated cardiomyocytes respectively in *Cbfβ*^Δ/Δ^ mice relative to control *Cbfβ*^f*l/fl*^ *mice* (Fig. 5f and 5g).

Echocardiography was performed the day before MI (day 0) and at 1 and 7 days following MI to assess LV function in *Cbfβ*^Δ/Δ^ mice relative to control *Cbfβ^fl/fl^* mice (Fig. 5h). As expected, at 1 day post-MI there was a decline in cardiac systolic function (measured by %FS) in *Cbfβ^fl/fl^ mice* which was maintained at a low level from day 1 to 7. By contrast, *Cbfβ*^Δ/Δ^ mice demonstrated a preserved contractile function at day 7 post-MI relative to control *Cbfβ*^f*l/fl*^ mice (35.2±2.2 *vs*. 25.3±1.5% FS; *P*<0.05, Fig. 5h).

These data support the findings with Ro5-3335 that limiting RUNX1 function by reducing availability of its essential co-factor *CBFβ* in cardiomyocytes preserves cardiac function.

### RNA-sequencing analyses of LV myocardium in *Cbfβ*^Δ/Δ^ and *Cbfβ*^fl/fl^ mice

We next determined whether there was any similarity between the pathways altered in *Cbfβ*^Δ/Δ^ mice to our findings with *Runx1*^Δ/Δ^ mice (Fig. 2) using an unbiased RNAseq approach. LV myocardium taken before MI (day 0) and 7 days after MI was compared in both *Cbfβ*^Δ/Δ^ and *Cbfβ*^fl/fl^ control mice. Using a FDR of 0.05, there were 7860 differentially expressed genes in *Cbfβ*^fl/fl^ control mice when comparing the day 0 and day 7 time points (Figure 6a). By contrast, there were only 2995 differentially expressed genes in *Cbfβ*^Δ/Δ^ mice (Fig. 6a). There were 2747 differentially expressed genes that were common to both *Cbfβ*^Δ/Δ^ and *Cbfβ*^fl/fl^ control mice leaving 5113 differentially expressed genes unique to *Cbfβ* ^fl/fl^ mice and not significantly affected in *Cbfβ*^Δ/Δ^ (Fig.6a and 6b). These overall numbers are very similar to those shown in Runx1^Δ/Δ^ mice (Fig. 2). IPA software was used to determine the enriched pathways and functions specific to the 5113 differentially expressed genes unique to *Cbfβ* ^fl/fl^ mice compared to the 248 differentially expressed genes unique to the *Cbfβ*^Δ/Δ^ mice. As with our finding with *Runx1*^Δ/Δ^ mice, oxidative phosphorylation was the most significantly changed pathway (Fig.6c).

**Figure 6.**
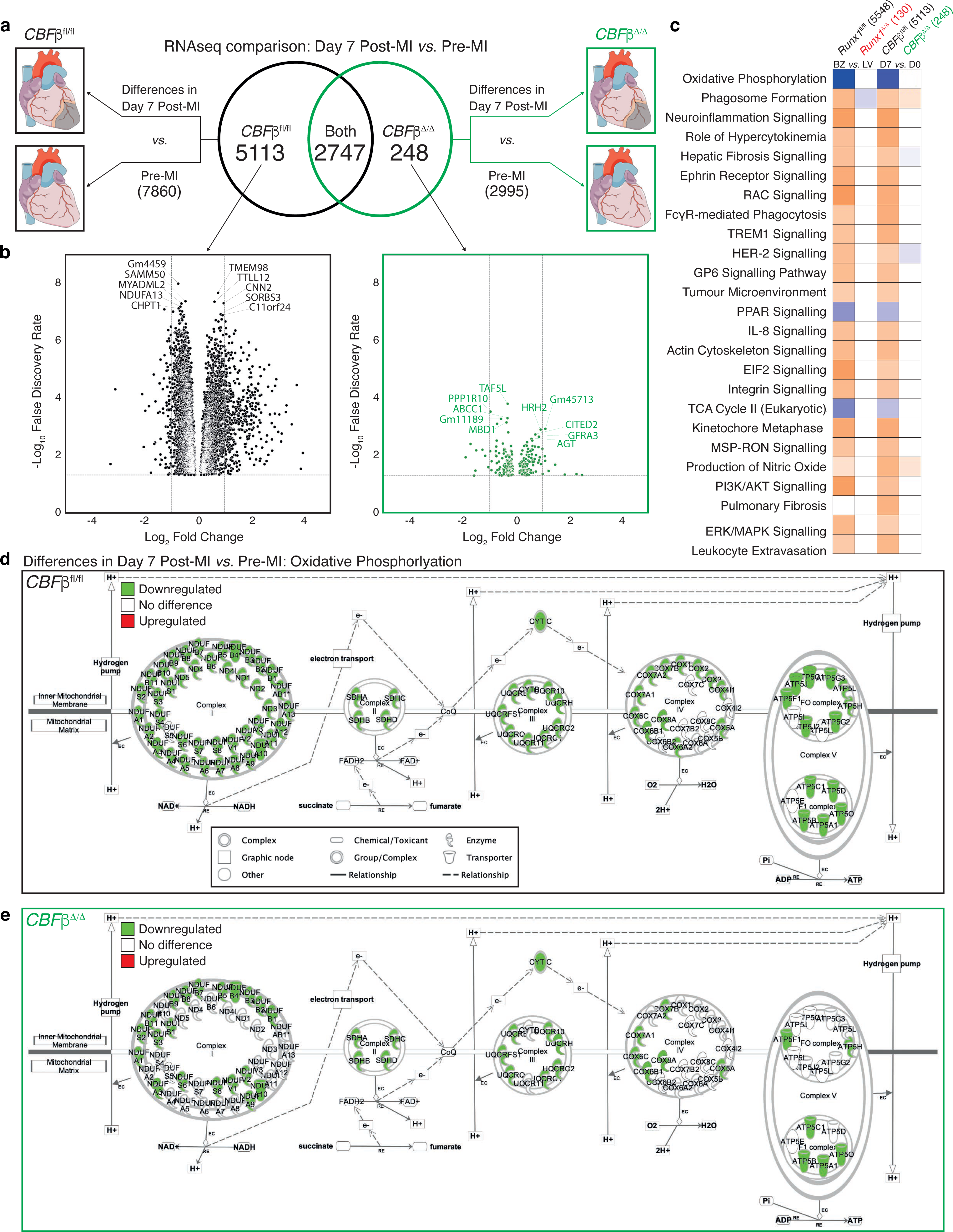
RNA sequencing comparisons and analysis between hearts 7 days post-myocardial infarction (MI) and pre-MI from *Cbfβ ^fl/fl^* and *Cbfβ* ^Δ/Δ^ mice. **(a)** Schematic of comparison and Venn diagram of gene expression. **(b)** Volcano plots of differentially regulated genes unique to *Cbfβ* ^fl/fl^ mice (left, black) and *Cbfβ* ^Δ/Δ^ mice (right, green) 7 days post-MI compared to pre-MI. Top five upregulated and down regulated genes based on false discovery rate (FDR) noted. **(c)** Enriched biological pathway from ingenuity pathway analysis (IPA) analysis generated from unique differences in Runx1^fl/fl^ (border zone, BZ *vs* remote zone, RZ) and *Cbfβ* ^fl/fl^ (post-MI *vs* Pre-MI) mice ranked on Z-score. Blue to orange heatmap and symbol colours represent predicted inhibition (blue) and activation (orange) or no change (white) of pathways based on Z-score. Schematic representation of genes involved in oxidative phosphorylation complexes from RNAseq IPA pathway analysis generated from all gene differences (downregulated – green, upregulated – red, no difference – white) between day 7 post-MI and pre-MI in **(d)** *Cbfβ* ^fl/fl^ (7860 differentially expressed genes) and **(e)** *Cbfβ* ^Δ/Δ^ mice (2995 differentially expressed genes).

Further interrogation of the pathway revealed marked downregulation of 87% of genes involved in oxidative phosphorylation across all five inner mitochondrial complexes in *Cbfβ*^fl/fl^ mice (Fig. 6d). By contrast, only 47% of mitochondrial genes were downregulated in the *Cbfβ*^Δ/Δ^ mice demonstrating that cardiomyocyte deficiency in *CBFβ* or *Runx1* preserves mitochondrial gene expression following MI (Fig. 6e).

## DISCUSSION

Our study reveals original and important insights into two key aspects of RUNX1 function. The first finding relates to the impact of the spatially restricted increase of Runx1 in BZ myocardium observed within the first day following MI. We demonstrate this phenomenon drives abnormal SR-mediated calcium handling, downregulation of mitochondrial gene expression, and a reduced number of mitochondria in the BZ relative to the RZ. Second, we demonstrate the therapeutic potential of inhibiting RUNX1 activity/expression using multiple translational approaches. These manipulations markedly and robustly preserve cardiac contractile function following MI. These discoveries have potential for clinical translation.

It is increasingly recognised that events that occur in the first few days post-MI play a critical role in the progression of LV remodelling during the following weeks and months^9^. LV contractile function is substantially reduced as early as 1-day post-MI^1–4^. Critically at this time point, across species, there is a regional heterogeneity of contractile function across the LV wall contributing to impaired global LV contractile function and poor patient prognosis^8–10^. The most severe contractile dysfunction in the surviving myocardium is in the BZ region (where *Runx1* expression is highest) and least dysfunction in the RZ region (where *Runx1* expression is lowest)^6, 11, 12^. Calcium handling within cardiomyocytes is one of the key determinants of cardiac contractile function. Here, we demonstrate that abnormal calcium handling in BZ cardiomyocytes occurs as early as 1 day post-MI. The abnormalities include a reduced calcium transient amplitude, SR calcium content and SERCA activity in BZ cardiomyocytes relative to cardiomyocytes isolated from the RZ region. These data are supported by previous studies that demonstrate that the BZ region has reduced SERCA expression, calcium transient amplitude and contraction at 3-7 days following MI^13, 35^. In contrast, the amplitude of calcium release, SR calcium content and SERCA activity was not significantly different between the BZ and RZ region in *Runx1-*deficient mice post-MI demonstrating homogeneous calcium handling across regions that is likely to contribute to the improved LV contractile function observed in *Runx1-* deficient mice at 1 day following MI. Our previous results at the later time point of 2 weeks post-MI demonstrated a role for Runx1 in determining SERCA-mediated calcium uptake, SR calcium content and calcium transient amplitude^20^. The current study provides new insight in that this function of Runx1 is evident in BZ cardiomyocytes as early as 1 day post-MI.

Our RNAseq study comparing the BZ and RZ in *Runx1-*deficient mice produced remarkable evidence for a new role for RUNX1 in reducing mitochondria function and number in the BZ following MI. Mitochondria are essential for the generation of vast amounts of energy required by the heart in the form of ATP. The mitochondria generates more than 95% of this ATP by oxidative phosphorylation^36^. Myocardial infarction is commonly characterised by changes in energy/substrate metabolism and decreased mitochondrial function with abnormalities of ATP production being more severe in the BZ relative to the RZ^17, 37^. This pattern, which occurs across species (rodents, sheep, pigs, zebrafish, and humans), coincides with a parallel reduction in the expression of genes associated with oxidative phosphorylation, reduced BZ contractile function and progression to HF^17, 38–42^. Our study provides new insight demonstrating that *Runx1* in the BZ drives a reduction in the expression of genes associated with mitochondrial oxidative phosphorylation as early as 1 day post-MI and likely contributes to impaired BZ contractile function. Our electron microscopy revealed that as early as 1 day following MI both mitochondria number and architecture is reduced/impaired respectively in the BZ. By contrast. the BZ of our *Runx1-*deficient mice had preserved expression of genes associated with mitochondrial oxidative phosphorylation relative to the RZ, and displayed a higher number of structurally normal mitochondria relative to control hearts post-MI. Mitochondrial size (and therefore potentially changes to mitochondrial fusion/fission) were not altered by Runx1. This new link between RUNX1 and mitochondrial function has profound implications for cardiac contractility and patient outcome and warrants future investigation. For example, the extent to which the expression of genes associated with oxidative phosphorylation in the BZ of *Runx1* deficient mice post-MI is regulated by effects of RUNX1 on mitochondria number (via altered balance of mitochondrial destruction/biogenesis), a multitude of genes or a key regulator remains an intriguing area to explore.

Given the potential of *Runx1* deficiency in cardiomyocyte-specific genetically modified mice to improve two key indices of BZ cardiomyocyte contractility (i.e., calcium handling and mitochondrial function) we tested whether targeting RUNX1 expression/activity in the BZ of wild type mice could enhance contractility post-MI. We first used two viral vector-mediated gene delivery approaches of *Runx1*-shRNA to knockdown *Runx1* within the BZ. Both of these approaches resulted in a marked preservation of LV contractile function post-MI. The level of improved contractility was equivalent to that observed in cardiomyocyte-specific *Runx1* deficient mice^20^. One unexpected difference between the two approaches was that, in contrast to direct BZ myocardial injection of Ad-*Runx1*-shRNA, intravenous injection of AAV9-*Runx1*-shRNA produced a small but statistically significant reduction of infarct size following MI. It is unclear why this difference in infarct size was only present when using AAV9-*Runx1*-shRNA but it is unlikely to explain the preservation of contractile function since direct myocardial BZ injection of Ad-*Runx1*-shRNA produced the same level of preserved LV contractility without a change in infarct size. This conclusion is also supported by our previous published work in cardiomyocyte-specific *Runx1* deficient mice where infarct size was not altered, yet cardiac contractility was preserved post-MI^20^.

Our next approach was to use Ro5-3335, a small molecule specific inhibitor of RUNX1^31^. Cardiac contractile function was preserved post-MI when Ro5-3335 was delivered either before or immediately after MI (the latter being a potential translational approach). Ro5-3335 is able to directly interact with RUNX1 and its cofactor CBFβ significantly reducing the transcriptional activity of RUNX1^31^. Ro5-3335 has previously been shown to improve pulmonary hypertension and retinal angiogenesis by altering vascular remodelling, endothelial to haemopoietic transition and pulmonary macrophage activity^43, 44^. Given the ability of Ro5-3335 to inhibit RUNX1-dependent processes outside the heart it was important to determine that the preservation in contractile function observed in Ro5-3335 treated mice was related to a direct effect on cardiomyocytes. To that end, we developed a tamoxifen-inducible cardiomyocyte-specific *Cbfβ* deficient mouse to limit Runx1 activity in cardiomyocytes in a similar way to Ro5-3335. *CBFβ* deficient mice also demonstrated preserved LV contractile function thus confirming (alongside our four other approaches to inhibit RUNX1 within the heart) that inhibiting RUNX1 within cardiomyocytes is a therapeutically tractable approach with translational potential to preserve cardiac contractile function following MI. Similar to our findings in *Runx1-*deficient mice, the *Cbfβ-*deficient mice had preserved expression levels of oxidative phosphorylation genes following MI, thus highlighting a common RUNX1-dependent mechanism observed in more than one of our approaches to inhibit RUNX1 activity in cardiomyocytes. Given that RUNX1 is a master transcription factor that controls multiple cellular processes, we anticipate potential for non-cardiomyocyte effects of small molecule based RUNX1 inhibition (e.g., vascular remodelling) that may contribute also to improved LV function. Indeed, using Runx1 as a multi-targeted approach to cardioprotection could be more beneficial than single-target based therapy to prevent adverse cardiac remodelling^45^.

Although the current study focuses on RUNX1 in the context of MI our findings have broader implications for RUNX1 biology in other tissues and diseases (cardiac and non-cardiac)^19^. Increased RUNX1 expression is now observed in hypertension-induced cardiac hypertrophy^46^, diabetes-induced heart dysfunction^47^ and dilated cardiomyopathy^22, 48^ all of which involve impaired energy production^49, 50^. The atria of patients with atrial fibrillation also demonstrate increased Runx1 expression^51^. Whether expression of Runx1 alters propensity for arrhythmias warrants future investigation. With regards to non-cardiac diseases, increased RUNX1 expression is a key contributor to retinal vascular dysfunction^44^, liver disease^52^, septic shock^53^ and kidney dysfunction^54^. In support of our work on the therapeutic potential of RUNX1 inhibitors in the setting of myocardial infarction, Ro5-3335 has been successfully used to improve both cardiac^55^ and non-cardiac related diseases^56^. It is becoming increasingly clear that RUNX1 inhibitors may represent a future therapy for a wide range of diseases including MI. Furthermore, given the importance of our novel data linking increased RUNX1 expression to mitochondrial number and oxidative phosphorylation, this study opens up the potential to explore the importance of RUNX1 in cellular processes/diseases where mitochondrial oxidative phosphorylation gene expression is demonstrated to be decreased; for example, mitochondrial disorders^57^ and age-associated organ dysfunction^58^.

In summary, the BZ myocardium is an important hypocontractile region of the heart post-MI the pathology of which influences progression to adverse cardiac remodelling and heart failure. We have demonstrated that RUNX1 drives pathological changes within BZ cardiomyocytes as early as 1 day post-MI; in particular a reduction in SR calcium release, mitochondrial number and expression of genes involved in oxidative phosphorylation necessary for ATP production. Inhibition of *Runx1* by gene therapeutic approaches or the use of a small molecular inhibitor improves LV cardiac contractile function and represents a new translational approach with the potential to treat patients with MI and limit progression to heart failure.

## ACKNOWLEDGMENTS

Funding was received from the British Heart Foundation (PG/18/933548 and RG/20/6/35095) to CML and MG (RG/19/11/34633) and the Wellcome Trust (204820/Z/16/Z) to TPM. EARZ is a German Research Foundation Emmy Noether Fellow (DFG #396913060) and a member of the German Research Foundation Collaborative Research Centre SFB1425 (DFG #422681845). Ali Elbassioni is funded by the Newton-Mosharafa Fund. CRUK core funding to the Beatson Institute (A31287) and to Karen Blyth (A29799). The authors thank Michael Dunne, Margaret Bell, and Catherine Hawksby at the University of Glasgow for their surgical, animal, and technical assistance; as well as Dr Martin Schorb and Dr Rachel Mellwig at EMBL Heidelberg Electron Microscopy Core Facility for access to and assistance with electron microscopy and tomography. We thank Douglas Strathdee, the Transgenic Technologies Lab, BSU and Histology lab at the Beatson Institute. We thank Vicky Heath for editorial assistance.

## AUTHOR CONTRIBUTIONS

**Tamara P Martin:** Conceptualization, Methodology, Validation, Formal analysis, Investigation, Writing-Original draft and Review and Editing, Supervision, Project administration; **Eilidh A MacDonald:** Conceptualization, Methodology, Validation, Formal analysis, Investigation, Writing-Original draft and Review and Editing, Visualization, Supervision, Project administration; **Ashley Bradley:** Conceptualization, Methodology, Validation, Formal analysis, Investigation, Writing-Review and Editing; **Holly Watson:** Methodology, Formal analysis, Investigation; **Priyanka Saxena:** Validation, Formal analysis, Investigation; **Eva Rog-Zielinska:**Methodology, Validation, Formal analysis, Investigation; **Simon Fisher:** Formal analysis; **Catriona Booth:** Formal analysis; **Morna Campbell:** Formal analysis; **Ali Ali Mohamed Elbassioni:** Formal analysis, Investigation; **Ohood Almuzaini:**Formal analysis, Investigation; **Pawel Herzyk:** Formal analysis; **Karen Blyth:** Writing-Review and Editing; **Colin Nixon:** Investigation, Resources; **Lorena Zentilin:** Resources; **Colin Berry:** Writing-Review and Editing; **Thomas Braun:** Conceptualization, Writing-Review and Editing; **Mauro Giacca:** Resources; Writing-Review and Editing**; Martin McBride:** Formal analysis, Investigation, Writing-Review and Editing, Supervision; **Stuart A Nicklin:** Supervision, Writing-Review and Editing, Funding acquisition; **Ewan Cameron:** Conceptualization, Supervision, Writing-Review and Editing, Funding acquisition; **Christopher M Loughrey:** Conceptualization, Formal analysis, Writing-Original draft and Review and Editing, Supervision, Project administration, Resources, Data Curation, Funding acquisition.

## METHODS

### Mice

The care and use of animals were in accordance with the UK Government Animals (Scientific Procedures) Act 1986. All animal procedures were approved by the University of Glasgow Animal Welfare and Ethical Review Body and licensed by the Home Office, UK (project licence no. P05FEIF82). Mice were housed on a 12/12 h light/ dark cycle and fed and watered *ad libitum.* All animals used were male mice aged 10-12 weeks of age (weight, 25-30 g) and were randomly assigned to experimental groups.

### Generation of cardiomyocyte-specific *Runx1* and *Cbfβ* deficient mice

*Runx1*^Δ/Δ^ mice were generated as previously described^20^. *Cbfβ*^fl/fl^ mice^33^ (The Jackson Laboratory, ME, USA) were crossed with mice expressing tamoxifen-inducible Cre recombinase (MerCreMer) under the control of the cardiac-specific αMHC (α-myosin heavy chain)^34^ to generate the relevant test (*Cbfβ* ^Δ/Δ^) and littermate (*Cbfβ*^fl/fl^) control groups.

### Coronary artery ligation

Thoracotomy and left anterior descending coronary artery permanent ligation surgeries were performed on C57BL/6J (Envigo), *Runx1-*deficient, *Cbfβ-*deficient and respective floxed control mice using previously published standard approaches (see Extended Methods)^59, 60^.

### Calcium measurements

Cardiomyocytes from C57BL/6J and *Runx1-*deficient mice were isolated according to region (see Extended Methods). BZ and remote RZ were loaded with a calcium-sensitive fluorophore (5.0 μmol/L Fura-4F AM, Invitrogen), and perfused during field stimulation (1.0 Hz, 2.0ms duration, stimulation voltage set to 1.5 times the threshold). Data were analysed offline as previously described^61^.

### RNA sequencing sample preparation and analysis

RNA was extracted using the miRNeasy Mini Kit (Qiagen, UK) from BZ and RZ myocardial tissue from control *Runx1*^fl/fl^ and *Runx1*^Δ/Δ^ mice pre-MI as well as 1 day post-MI and whole LV myocardial tissue from *Cbfβ*^fl/fl^ and *Cbfβ*^Δ/Δ^ pre-MI and 7 days post-MI. Sequencing libraries were enriched using polyA tail selection and samples were run on a Next Seq500 Sequencing system (Illumina) at the Glasgow Polyomics research facility (see Extended Methods). Global functional, network, and canonical pathway analyses were performed using Ingenuity Pathways Analysis (IPA). Genes shown in the article text and figures were generated by IPA and therefore might exclude lncRNAs and pseudogenes that are not recognised by the IPA software.

### Electron Microscopy

Hearts were perfused with cardioplegic solution, immediately followed by perfusion-fixation with iso-osmotic Karnovsky’s fixative (2.4% sodium cacodylate, 0.75% paraformaldehyde, 0.75% glutaraldehyde) and then separated into BZ and RZ. Thin (90 nm) and semi-thick (300 nm, for dual-axis electron tomography) sections were prepared and imaged at the Electron Microscopy Core Facility, European Molecular Biology Laboratory (EMBL) Heidelberg, using 300 kV Tecnai TF30 (FEI Company, now Thermo-Fisher Scientific, Eindhoven, The Netherlands). Tilt series were aligned, reconstructed, and combined using IMOD. Mitochondrial density, quantity, and damage was analysed using IMOD software and ImageJ (see Extended Methods).

### Adenoviral knockdown of *Runx1* in the border zone region

The Ad-*Runx1*-shRNA^62^ and a random scramble sequence (Ad-scramble-shRNA) were purchased from Vector Biolabs (USA; Gene ID: 12394). Following coronary artery ligation, the BZ area was injected with 5 x 10 μL of either Ad-scramble-shRNA or Ad-*Runx1*-shRNA (1 x10 ^9^ viral particles per heart) immediately following MI (see Extended Methods).

### AAV-mediated knockdown of *Runx1*

Following coronary artery ligation, mice were intravenously injected via the tail vein with either AAV9-scramble-shRNA or AAV9-*Runx1*-shRNA (1x10^11^ virus particles per mouse) in a randomised fashion immediately following MI (see Extended Methods).

### Ro5-3335 knockdown of *Runx1*

C57BL/6J mice were randomly assigned to receive either vehicle (100% DMSO) or 20 mg/ kg of Ro5-3335^43^ (Tocris-Bioscience, UK) given subcutaneously. Mice assigned to protocol 1 received either Ro5-3335 or vehicle at 7, 5, 3 and 1 day before coronary artery ligation then again at 1, 3, 5 and 7 days following coronary artery ligation. Mice assigned to protocol 2 received Ro5-3335 or vehicle at the time of coronary artery ligation and on consecutive days until day 7.

### Echocardiography

M-mode measurements were performed prior to left anterior descending coronary artery ligation and in the days following as previously published^20^.

### Histology

Quantification of regional areas and infarct size was performed on Picrosirius Red-stained histological sections with ImageJ and Adobe Photoshop as previously described^20^. RNAscope with probes to specifically identify cardiomyocyte nuclei (pericentriolar material 1; *PCM1*) and *Runx1* was performed. For each heart, positive (*Ppib* and *Polr2a*) and negative controls (bacterial dapB) were run (Extended figure 4).

### Statistics

Data were expressed as mean ± SEM. Comparisons between two experimental groups were performed with the Student’s t-test on raw data before normalization to percentage change where appropriate. Comparisons between more than two groups were conducted on raw data with ANOVA. In experiments where multiple isolated cardiomyocyte observations (*n*) were obtained from each heart, we first ensured normality of the data distribution and then determined the differences between control and experimental mice using mean data from each heart (and not individual cardiomyocytes; IBM SPSS Statistics, version 22).

## REFERENCES

1. Sandmann S, Min JY, Meissner A and Unger T. Effects of the calcium channel antagonist mibefradil on haemodynamic parameters and myocardial Ca(^2+^)-handling in infarct-induced heart failure in rats. Cardiovascular research. 1999;44:67–80.

2. Yue P, Long CS, Austin R, Chang KC, Simpson PC and Massie BM. Post-infarction heart failure in the rat is associated with distinct alterations in cardiac myocyte molecular phenotype. Journal of molecular and cellular cardiology. 1998;30:1615–30.

3. Palojoki E, Saraste A, Eriksson A, Pulkki K, Kallajoki M, Voipio-Pulkki LM and Tikkanen I. Cardiomyocyte apoptosis and ventricular remodeling after myocardial infarction in rats. American journal of physiology Heart and circulatory physiology. 2001;280:H2726–31.

4. Song K, Nam YJ, Luo X, Qi X, Tan W, Huang GN, Acharya A, Smith CL, Tallquist MD, Neilson EG, Hill JA, Bassel-Duby R and Olson EN. Heart repair by reprogramming non-myocytes with cardiac transcription factors. Nature. 2012;485:599–604.

5. Konstam MA, Kramer DG, Patel AR, Maron MS and Udelson JE. Left ventricular remodeling in heart failure: current concepts in clinical significance and assessment. JACC Cardiovasc Imaging. 2011;4:98–108.

6. Kramer CM, Lima JA, Reichek N, Ferrari VA, Llaneras MR, Palmon LC, Yeh IT, Tallant B and Axel L. Regional differences in function within noninfarcted myocardium during left ventricular remodeling. Circulation. 1993;88:1279–88.

7. Roger VL. Epidemiology of heart failure. Circulation research. 2013;113:646–659.

8. Jackson BM, Gorman JH, Moainie SL, Guy TS, Narula N, Narula J, John-Sutton MG, Edmunds LH, Jr. and Gorman RC. Extension of borderzone myocardium in postinfarction dilated cardiomyopathy. J AmCollCardiol. 2002;40:1160–1167.

9. French BA and Kramer CM. Mechanisms of Post-Infarct Left Ventricular Remodeling. Drug Discov Today Dis Mech. 2007;4:185–196.

10. Heidary S, Patel H, Chung J, Yokota H, Gupta SN, Bennett MV, Katikireddy C, Nguyen P, Pauly JM, Terashima M, McConnell MV and Yang PC. Quantitative tissue characterization of infarct core and border zone in patients with ischemic cardiomyopathy by magnetic resonance is associated with future cardiovascular events. Journal of the American College of Cardiology. 2010;55:2762–8.

11. Epstein FH, Yang Z, Gilson WD, Berr SS, Kramer CM and French BA. MR tagging early after myocardial infarction in mice demonstrates contractile dysfunction in adjacent and remote regions. Magnetic resonance in medicine : official journal of the Society of Magnetic Resonance in Medicine / Society of Magnetic Resonance in Medicine. 2002;48:399–403.

12. Kramer CM, Rogers WJ, Theobald TM, Power TP, Petruolo S and Reichek N. Remote noninfarcted region dysfunction soon after first anterior myocardial infarction. A magnetic resonance tagging study. Circulation. 1996;94:660–6.

13. Yoshiyama M, Takeuchi K, Hanatani A, Kim S, Omura T, Toda I, Teragaki M, Akioka K, Iwao H and Yoshikawa J. Differences in expression of sarcoplasmic reticulum Ca^2+^-ATPase and Na^+^-Ca^2+^ exchanger genes between adjacent and remote noninfarcted myocardium after myocardial infarction. Journal of molecular and cellular cardiology. 1997;29:255–64.

14. Mork HK, Sjaastad I, Sande JB, Periasamy M, Sejersted OM and Louch WE. Increased cardiomyocyte function and Ca^2+^ transients in mice during early congestive heart failure. Journal of molecular and cellular cardiology. 2007;43:177–86.

15. Mork HK, Sjaastad I, Sejersted OM and Louch WE. Slowing of cardiomyocyte Ca^2+^ release and contraction during heart failure progression in postinfarction mice. American journal of physiology Heart and circulatory physiology. 2009;296:H1069–79.

16. Unsold B, Kaul A, Sbroggio M, Schubert C, Regitz-Zagrosek V, Brancaccio M, Damilano F, Hirsch E, Van Bilsen M, Munts C, Sipido K, Bito V, Detre E, Wagner NM, Schafer K, Seidler T, Vogt J, Neef S, Bleckmann A, Maier LS, Balligand JL, Bouzin C, Ventura-Clapier R, Garnier A, Eschenhagen T, El-Armouche A, Knoll R, Tarone G and Hasenfuss G. Melusin protects from cardiac rupture and improves functional remodelling after myocardial infarction. Cardiovascular research. 2014;101:97–107.

17. van Duijvenboden K, de Bakker DEM, Man JCK, Janssen R, Gunthel M, Hill MC, Hooijkaas IB, van der Made I, van der Kraak PH, Vink A, Creemers EE, Martin JF, Barnett P, Bakkers J and Christoffels VM. Conserved NPPB+ Border Zone Switches from MEF2 to AP-1 Driven Gene Program. Circulation. 2019;140(10):864–879.

18. Blyth K, Cameron ER and Neil JC. The RUNX genes: gain or loss of function in cancer.NatRevCancer. 2005;5:376–387.

19. Riddell A, McBride M, Braun T, Nicklin SA, Cameron E, Loughrey CM and Martin TP. RUNX1: an emerging therapeutic target for cardiovascular disease. Cardiovascular research. 2020;116:1410–1423.

20. McCarroll CS, He W, Foote K, Bradley A, McGlynn K, Vidler F, Nixon C, Nather K, Fattah C, Riddell A, Bowman P, Elliott EB, Bell M, Hawksby C, MacKenzie SM, Morrison LJ, Terry A, Blyth K, Smith GL, McBride MW, Kubin T, Braun T, Nicklin SA, Cameron ER and Loughrey CM. Runx1 Deficiency Protects Against Adverse Cardiac Remodeling After Myocardial Infarction. Circulation. 2018;137:57–70.

21. Gattenlohner S, Waller C, Ertl G, Bultmann BD, Muller-Hermelink HK and Marx A. NCAM(CD56) and RUNX1(AML1) are up-regulated in human ischemic cardiomyopathy and a rat model of chronic cardiac ischemia. AmJ Pathol. 2003;163:1081–1090.

22. Kubin T, Poling J, Kostin S, Gajawada P, Hein S, Rees W, Wietelmann A, Tanaka M, Lorchner H, Schimanski S, Szibor M, Warnecke H and Braun T. Oncostatin M is a major mediator of cardiomyocyte dedifferentiation and remodeling. Cell stem cell. 2011;9:420–32.

23. Trafford AW, Diaz ME and Eisner DA. Anovel ,rapid and reversible method to measure calcium buffering and time course of total sarcoplasmic reticulum calcium content in cardiac ventricular myocytes. Pflugers Arch - Eur J Physiol. 1999;437:501–503.

24. Bassani JW, Yuan W and Bers DM. Fractional SR Ca^2+^ release is regulated by trigger Ca^2+^ and SR Ca^2+^ content in cardiac myocytes. AmJPhysiol. 1995;268:C1313–C1319.

25. Eisner DA, Choi HS, Diaz ME, O’Neill SC and Trafford AW. Intergrative analysis of calcium cycling in cardiac muscle. Circulation research. 2000;87:1087–1094.

26. Diaz ME, Graham HK and Trafford AW. Enhanced sarcolemmal Ca(^2+^) efflux reduces sarcoplasmic reticulum Ca(^2+^) content and systolic Ca(^2+^) in cardiac hypertrophy. Cardiovascular research. 2004;62:538–547.

27. Piacentino V, III, Weber CR, Chen X, Weisser-Thomas J, Margulies KB, Bers DM and Houser SR. Cellular basis of abnormal calcium transients of failing human ventricular myocytes. CircRes. 2003;92:651–658.

28. Cannata A, Ali H, Sinagra G and Giacca M. Gene Therapy for the Heart Lessons Learned and Future Perspectives. Circulation research. 2020;126:1394–1414.

29. Mendell JR, Al-Zaidy S, Shell R, Arnold WD, Rodino-Klapac LR, Prior TW, Lowes L, Alfano L, Berry K, Church K, Kissel JT, Nagendran S, L’Italien J, Sproule DM, Wells C, Cardenas JA, Heitzer MD, Kaspar A, Corcoran S, Braun L, Likhite S, Miranda C, Meyer K, Foust KD, Burghes AHM and Kaspar BK. Single-Dose Gene-Replacement Therapy for Spinal Muscular Atrophy. N Engl J Med. 2017;377:1713–1722.

30. Fattah C, Nather K, McCarroll CS, Hortigon-Vinagre MP, Zamora V, Flores-Munoz M, McArthur L, Zentilin L, Giacca M, Touyz RM, Smith GL, Loughrey CM and Nicklin SA. Gene Therapy With Angiotensin-(1-9) Preserves Left Ventricular Systolic Function After Myocardial Infarction. Journal of the American College of Cardiology. 2016;68:2652–2666.

31. Cunningham L, Finckbeiner S, Hyde RK, Southall N, Marugan J, Yedavalli VR, Dehdashti SJ, Reinhold WC, Alemu L, Zhao L, Yeh JR, Sood R, Pommier Y, Austin CP, Jeang KT, Zheng W and Liu P. Identification of benzodiazepine Ro5-3335 as an inhibitor of CBF leukemia through quantitative high throughput screen against RUNX1-CBFbeta interaction. Proc Natl Acad Sci U S A. 2012;109:14592–7.

32. Whitmore HAB, Amarnani D, O’Hare M, Delgado-Tirado S, Gonzalez-Buendia L, An M, Pedron J, Bushweller JH, Arboleda-Velasquez JF and Kim LA. TNF-alpha signaling regulates RUNX1 function in endothelial cells. FASEB J. 2021;35:e21155.

33. Naoe Y, Setoguchi R, Akiyama K, Muroi S, Kuroda M, Hatam F, Littman DR and Taniuchi I. Repression of interleukin-4 in T helper type 1 cells by Runx/Cbf beta binding to the Il4 silencer. J Exp Med. 2007;204:1749–55.

34. Sohal DS, Nghiem M, Crackower MA, Witt SA, Kimball TR, Tymitz KM, Penninger JM and Molkentin JD. Temporally regulated and tissue-specific gene manipulations in the adult and embryonic heart using a tamoxifen-inducible Cre protein. CircRes. 2001;89:20–25.

35. Licata A, Aggarwal R, Robinson RB and Boyden P. Frequency dependent effects on calcium transients and cell shortening in myocytes that survive in the infarcted heart. Cardiovascular research. 1997;33:341–50.

36. Maack C and O’Rourke B. Excitation-contraction coupling and mitochondrial energetics. Basic research in cardiology. 2007;102:369–92.

37. Ait-Aissa K, Blaszak SC, Beutner G, Tsaih SW, Morgan G, Santos JH, Flister MJ, Joyce DL, Camara AKS, Gutterman DD, Donato AJ, Porter GA, Jr. and Beyer AM. Mitochondrial Oxidative Phosphorylation defect in the Heart of Subjects with Coronary Artery Disease. Scientific reports. 2019;9:7623.

38. Hu Q, Wang X, Lee J, Mansoor A, Liu J, Zeng L, Swingen C, Zhang G, Feygin J, Ochiai K, Bransford TL, From AH, Bache RJ and Zhang J. Profound bioenergetic abnormalities in peri-infarct myocardial regions. AmJ Physiol Heart CircPhysiol. 2006;291:H648–H657.

39. Honkoop H, de Bakker DE, Aharonov A, Kruse F, Shakked A, Nguyen PD, de Heus C, Garric L, Muraro MJ, Shoffner A, Tessadori F, Peterson JC, Noort W, Bertozzi A, Weidinger G, Posthuma G, Grun D, van der Laarse WJ, Klumperman J, Jaspers RT, Poss KD, van Oudenaarden A, Tzahor E and Bakkers J. Single-cell analysis uncovers that metabolic reprogramming by ErbB2 signaling is essential for cardiomyocyte proliferation in the regenerating heart. Elife. 2019;8:e50163.

40. Wang J, He J, Fan Y, Xu F, Liu Q, He R and Yan R. Extensive mitochondrial proteome disturbance occurs during the early stages of acute myocardial ischemia. Exp Ther Med. 2022;23:85.

41. Lock MC, Tellam RL, Darby JRT, Soo JY, Brooks DA, Macgowan CK, Selvanayagam JB, Porrello ER, Seed M, Keller-Wood M and Morrison JL. Differential gene responses 3 days following infarction in the fetal and adolescent sheep heart. Physiological genomics. 2020;52:143–159.

42. Zimmermann M, Beer L, Ullrich R, Lukovic D, Simader E, Traxler D, Wagner T, Nemec L, Altenburger L, Zuckermann A, Gyongyosi M, Ankersmit HJ and Mildner M. Analysis of region specific gene expression patterns in the heart and systemic responses after experimental myocardial ischemia. Oncotarget. 2017;8:60809–60825.

43. Jeong EM, Pereira M, So EY, Wu KQ, Del Tatto M, Wen S, Dooner MS, Dubielecka PM, Reginato AM, Ventetuolo CE, Quesenberry PJ, Klinger JR and Liang OD. Targeting RUNX1 as a novel treatment modality for pulmonary arterial hypertension. Cardiovascular research 2022 (online ahead of print 9 January), doi:10.1093/cvr/cvac001.

44. Lam JD, Oh DJ, Wong LL, Amarnani D, Park-Windhol C, Sanchez AV, Cardona-Velez J, McGuone D, Stemmer-Rachamimov AO, Eliott D, Bielenberg DR, van Zyl T, Shen L, Gai X, D’Amore PA, Kim LA and Arboleda-Velasquez JF. Identification of RUNX1 as a Mediator of Aberrant Retinal Angiogenesis. Diabetes. 2017;66:1950–1956.

45. Davidson SM, Ferdinandy P, Andreadou I, Botker HE, Heusch G, Ibanez B, Ovize M, Schulz R, Yellon DM, Hausenloy DJ, Garcia-Dorado D and Action CC. Multitarget Strategies to Reduce Myocardial Ischemia/Reperfusion Injury: JACC Review Topic of the Week. Journal of the American College of Cardiology. 2019;73:89–99.

46. Ikeda S, Mizushima W, Sciarretta S, Abdellatif M, Zhai P, Mukai R, Fefelova N, Oka SI, Nakamura M, Del Re DP, Farrance I, Park JY, Tian B, Xie LH, Kumar M, Hsu CP, Sadayappan S, Shimokawa H, Lim DS and Sadoshima J. Hippo Deficiency Leads to Cardiac Dysfunction Accompanied by Cardiomyocyte Dedifferentiation During Pressure Overload. Circulation research. 2019;124:292–305.

47. Zhang X, Ma S, Zhang R, Li S, Zhu D, Han D, Li X, Li C, Yan W, Sun D, Xu B, Wang Y and Cao F. Oncostatin M-induced cardiomyocyte dedifferentiation regulates the progression of diabetic cardiomyopathy through B-Raf/Mek/Erk signaling pathway. Acta Biochim Biophys Sin (Shanghai). 2016;48:257–65.

48. Burke MA, Chang S, Wakimoto H, Gorham JM, Conner DA, Christodoulou DC, Parfenov MG, DePalma SR, Eminaga S, Konno T, Seidman JG and Seidman CE. Molecular profiling of dilated cardiomyopathy that progresses to heart failure. JCI Insight. 2016;1:e86898.

49. Dillmann WH. Diabetic Cardiomyopathy. Circulation research. 2019;124:1160–1162.

50. Rosca MG, Vazquez EJ, Kerner J, Parland W, Chandler MP, Stanley W, Sabbah HN and Hoppel CL. Cardiac mitochondria in heart failure: decrease in respirasomes and oxidative phosphorylation. Cardiovascular research. 2008;80:30–9.

51. Zhao G, Zhou J, Gao J, Liu Y, Gu S, Zhang X and Su P. Genome-wide DNA methylation analysis in permanent atrial fibrillation. Mol Med Rep. 2017;16:5505–5514.

52. Kaur S, Rawal P, Siddiqui H, Rohilla S, Sharma S, Tripathi DM, Baweja S, Hassan M, Vlaic S, Guthke R, Thomas M, Dayoub R, Bihari C, Sarin SK and Weiss TS. Increased Expression of RUNX1 in Liver Correlates with NASH Activity Score in Patients with Non-Alcoholic Steatohepatitis (NASH). Cells. 2019;8.

53. Luo MC, Zhou SY, Feng DY, Xiao J, Li WY, Xu CD, Wang HY and Zhou T. Runt-related Transcription Factor 1 (RUNX1) Binds to p50 in Macrophages and Enhances TLR4-triggered Inflammation and Septic Shock. J Biol Chem. 2016;291:22011–22020.

54. Formica C, Malas T, Balog J, Verburg L, t Hoen PAC and Peters DJM. Characterisation of transcription factor profiles in polycystic kidney disease (PKD): identification and validation of STAT3 and RUNX1 in the injury/repair response and PKD progression. J Mol Med (Berl). 2019;97:1643–1656.

55. Zhang D, Liang C, Li P, Yang L, Hao Z, Kong L, Tian X, Guo C, Dong J, Zhang Y and Du B. Runt-related transcription factor 1 (Runx1) aggravates pathological cardiac hypertrophy by promoting p53 expression. J Cell Mol Med. 2021;25:7867–7877.

56. Delgado-Tirado S, Amarnani D, Zhao G, Rossin EJ, Eliott D, Miller JB, Greene WA, Ramos L, Arevalo-Alquichire S, Leyton-Cifuentes D, Gonzalez-Buendia L, Isaacs-Bernal D, Whitmore HAB, Chmielewska N, Duffy BV, Kim E, Wang HC, Ruiz-Moreno JM, Kim LA and Arboleda-Velasquez JF. Topical delivery of a small molecule RUNX1 transcription factor inhibitor for the treatment of proliferative vitreoretinopathy. Scientific reports. 2020;10:20554.

57. Fernandez-Vizarra E and Zeviani M. Mitochondrial disorders of the OXPHOS system. FEBS letters. 2021;595:1062–1106.

58. Amorim JA, Coppotelli G, Rolo AP, Palmeira CM, Ross JM and Sinclair DA. Mitochondrial and metabolic dysfunction in ageing and age-related diseases. Nature Reviews Endocrinology. 2022.

59. Martin TP, MacDonald EA, Elbassioni AAM, O’Toole D, Zaeri AAI, Nicklin SA, Gray GA and Loughrey CM. Preclinical models of myocardial infarction: from mechanism to translation. Br J Pharmacol. 2021.

60. He W, McCarroll CS, Nather K, Ford K, Mangion K, Riddell A, O’Toole D, Zaeri A, Corcoran D, Carrick D, Lee MMY, McEntegart M, Davie A, Good R, Lindsay MM, Eteiba H, Rocchiccioli P, Watkins S, Hood S, Shaukat A, McArthur L, Elliott EB, McClure J, Hawksby C, Martin T, Petrie MC, Oldroyd KG, Smith GL, Oxford Acute Myocardial Infarction S, Channon KM, Berry C, Nicklin SA and Loughrey CM. Inhibition of myocardial cathepsin-L release during reperfusion following myocardial infarction improves cardiac function and reduces infarct size. Cardiovascular research. 2021 (online ahead of print 16 June), doi:10.1093/cvr/cvab204.

61. Elliott EB, Hasumi H, Otani N, Matsuda T, Matsuda R, Kaneko N, Smith GL and Loughrey CM. K201 (JTV-519) alters the spatiotemporal properties of diastolic Ca(^2+^) release and the associated diastolic contraction during beta-adrenergic stimulation in rat ventricular cardiomyocytes. Basic Res Cardiol. 2011;106(6):1009–22.

62. Wang J, Wang X, Holz JD, Rutkowski T, Wang Y, Zhu Z and Dong Y. Runx1 is critical for PTH-induced onset of mesenchymal progenitor cell chondrogenic differentiation. PloS one. 2013;8:e74255.

