## extended for "Inhibiting Runx1 protects heart function after myocardial infarction"

**Extended Figure 1. Diastolic calcium transient in *Runx1*$^{\Delta/\Delta}$ mice and C57BL/6J mice 1 day post myocardial infarction (MI). (a)** Diastolic calcium ($Ca^{2+}$) transient before MI in C57BL6 mice in the remote zone (RZ) *n*=76 (7 hearts) and border zone (BZ) *n*=43 (hearts). **(b)** Diastolic $Ca^{2+}$ transient at 1 day post-MI in C57BL6 mice in the RZ *n*=64 (9 hearts) and BZ *n*=30 (9 hearts). **(c)** Diastolic $Ca^{2+}$ transient at 1 day post-MI in *Runx1*$^{\Delta/\Delta}$ mice in the RZ *n*=16 (4 hearts) and BZ *n*=9 (4 hearts). Error bars represent mean ± SEM. Results not statistically significant, linear mixed modelling.

**Extended Figure 2. Genes involved in oxidative phosphorylation from RNAseq Ingenuity pathway analysis (IPA).** List of genes downregulated (green) or upregulated (red) in oxidative phosphorylation complexes from RNAseq IPA pathway analysis for each group. There were no oxidative phosphorylation (OxPhos) genes altered in the border zone *vs.* remote zone of *Runx1*$^{\Delta/\Delta}$ mice, so all changes were unique to *Runx1*$^{fl/fl}$. Between Day 7 post-MI and pre-MI there were changes in *CBFβ*$^{\Delta/\Delta}$ mice (turquoise box), however all changes were shared with *CBFβ*$^{fl/fl}$. There were also genes downregulated in *CBFβ*$^{fl/fl}$ mice but not *CBFβ*$^{\Delta/\Delta}$ mice (i.e. unique changes), all genes changed in *CBFβ*$^{fl/fl}$ mice (shared and unique) are contained within the yellow box.

**Extended Figure 3. *Runx1* mRNA expression in IP1B cells transduced with Ad-scramble-shRNA or Ad-*Runx1*-shRNA.** *Runx1* mRNA expression relative to *Peptidylprolyl Isomerase B* measured by real-time quantitative polymerase chain reaction in IP1B cells transduced with Ad-*Runx1*-shRNA (*n*=5; biological replicates) *vs*. Ad-scramble-shRNA (*n*=5; biological replicates) at (**a**) 500 PFU/cell or (**b**) 1000 PFU/cell. Error bars represent mean ± SEM. **P*<0.05, Student *t* test.

**Extended Figure 4. Control RNAscope images and cardiomyocyte and non-cardiomyocyte expression using Pericentriolar material 1 (*PCM-1*).** RNA in situ hybridisation using RNAscope at 7 days post-myocardial infarction (MI) in Ad-*Runx1*-shRNA (*n*=4) *vs.* Ad-scramble-shRNA (*n*=4) and at 7 days post-MI in AAV-*Runx1*-shRNA (*n*=3) *vs.* AAV-scramble-shRNA (*n*=3). (**a**) Mean quantification of *PCM-1* positive (cardiomyocyte) nuclei expressed as the percentage of total nuclei in Ad-*Runx1*/scramble-shRNA mice. (**b**) Mean quantification of *PCM-1* negative (non-cardiomyocyte) nuclei expressed as a percentage of total nuclei Ad-*Runx1*/scramble-shRNA mice. (**c**) Mean quantification of

*PCM-1* positive (cardiomyocyte) nuclei expressed as the percentage of total nuclei in Ad-*Runx1*/scramble-shRNA mice. (**d**) Mean quantification of *PCM-1* negative (non-cardiomyocyte) nuclei expressed as the percentage of total nuclei Ad-*Runx1*/scramble-shRNA mice. Error bars represent mean ±SEM. (**e**) positive control slides using *Peptidylprolyl Isomerase B (*pink puncate dots*)* and *Polr2 (*brown punctate dots*)* probes in Ad-scramble-shRNA MI mice and (**f**) positive *Ppib* and *Polr2* probes in Ad-*Runx1*-shRNA MI mice. (**g**) negative controls using bacterial bacillus subtilis dihydodipicolinate reductase (*dapB*).

**Extended Table 1. Raw data quantifying calcium ($Ca^{2+}$) handling parameters.** $Ca^{2+}$ transient peak (nmol.$L^{-1}$), $Ca^{2+}$ transient minimum (nmol.$L^{-1}$), $Ca^{2+}$ transient amplitude (nmol.$L^{-1}$), caffeine-induced $Ca^{2+}$ transient amplitude (nmol.$L^{-1}$), SERCA activity ($K_{SERCA,}$ $S^{-1}$), time constant of caffeine-induced transient decay (NCX activity, $S^{-1}$) and calcium transient peak to minimum ratio in C56BL/6J mice before MI in the remote zone (RZ; *n*=76 cardiomyocytes, *n*=9 hearts) and border zone (BZ; *n*=43 cardiomyocytes, n=9 hearts) and after MI in the RZ (*n*=64 cardiomyocytes, *n*=7 hearts) and BZ (*n*=30 cardiomyocytes, *n*=7 hearts) and at 1-day post-MI in the RZ (*n*=16 cardiomyocytes, *n*=4 hearts) and BZ (*n*=9 cardiomyocytes, *n*=4 hearts) of *Runx1*-deficient mice and in the RZ (*n*=20 cardiomyocytes, *n*=5 hearts) and BZ (*n*=31 cardiomyocytes, *n*=5 hearts) of $Runx1^{fl/fl}$ mice. *$P$<0.05, Student *t* test.

**Extended Table 2. Raw Echocardiography parameters.** Fractional shortening (FS%), left ventricular diameter at end diastole (LIVDd, mm), left ventricular diameter at end systole (LVIDs, mm), left ventricular posterior wall thickness at end diastole (LVPWd, mm) and left ventricular posterior wall thickness at end systole (LVPWs, mm) in Ad-*Runx1*-shRNA myocardial infarction (MI) mice (*n*=8; day 0, *n*=8; day 1, *n*=7; day 2, *n*=7; day 7) *vs.* Ad-scramble-shRNA MI mice (n=8; day 0, *n*=8; day 1, *n*=7; day 2, *n*=7; day 7). FS, LVIDd, LVIDs, LVPWd and LVPWs in AAV-*Runx1*-shRNA MI mice (*n*=9; day 0, *n*=9; day 1, *n*=9; day 7) vs. AAV-scramble-shRNA MI mice (*n*=10; day 0, *n*=10; day 1, *n*=9; day 7). FS, LVIDd, LVIDs, LVPWd and LVPWs in Ro5-3335 MI mice (Protocol 1: *n*=6; day 0, *n*=6; day 1, *n*=5; day 3, *n*=5; day 7. Protocol 2: *n*=8; day 0, *n*=8; day 1, *n*=6; day 3, *n*=6; day 7) *vs.* vehicle (DMSO) MI mice (Protocol 1: *n*=6; day 0, *n*=5; day 1, *n*=3; day 3, *n*=4; day 7. Protocol 2: *n*=7; day 0, *n*=8; day 1, *n*=3; day 3, *n*=7; day 7). FS, LVIDd, LVIDs, LVPWd and LVPWs in *Cbfβ*$^{Δ/Δ}$ MI mice (*n*=8; day 0, *n*=6; day 1, *n*=6; day 7) vs. *Cbfβ* $^{fl/fl}$ MI mice (*n*=6; day 0, *n*=;5 day 1, *n*=6; day 7).

#### EXTENDED METHODS

##### Coronary artery ligation

Thoracotomy and left anterior descending coronary artery permanent ligation were performed on C57BL/6J (Envigo), *Runx1*-deficient, *Cbfβ*-deficient and respective floxed control mice aged 10-12 weeks (weight 25–30 g). Mice were initially anaesthetised by 4% isoflourane (Isoflo, Abbott Laboratories, USA) receiving oxygen at 1 L/min. Mice received preoperative analgesia of 5 mg/kg carprofen (Rimadyl; Pfizer Animal Health, UK) and 0.1 mg/kg buprenorphine (Vetergesic; Reckitt Benkiser Healthcare Ltd, UK) delivered subcutaneously in a single injection made up to 0.4 mL with saline. Mice were endotracheally intubated and mechanically ventilated at 125 breath/min with a tidal volume of 120 µL. Isofluorane was reduced to 3% initially and then gradually reduced throughout the procedure. A 1 cm skin incision was made perpendicular to the sternum and across the rib cage. The skin and thoracic muscles were carefully retracted back, and the intercostal muscles blunt dissected. The pericardial sac was opened and removed to expose the heart and provide access for ligation. The left anterior descending (LAD) coronary artery was ligated with 9-0 non-adsorbable nylon (W2829 Ethilon; Johnson & Johnson, UK) 1.5 mm distal to the left atrial appendage. Three 6-0 non-absorbable prolene pre-placed sutures (W8711; Johnson & Johnson, UK) were inserted around the ribs and the lungs re-inflated. The thoracic muscles were returned to original positions and the skin closed with interrupted stitches using 6-0 absorbable vicryl (W9575; Johnson & Johnson, UK).

##### Calcium measurements

Cardiomyocytes from C57BL/6J and *Runx1*-deficient mice were isolated as previously described[1]. To isolate cells according to region, cardiomyocytes were dissociated from a 2mm strip of BZ tissue adjacent to the IZ. After the BZ, a 2mm strip of the LV was removed and discarded. Cardiomyocytes were dissociated from the remaining LV tissue (the remote LV; Figure 1a). The Fura-4F fluorescence ratio (340/380- nm excitation) was measured with a spinning wheel spectrophotometer (Cairn

Research Ltd; sampling rate of 5.0 kHz) to measure the cardiomyocyte intracellular calcium concentration ($[Ca^{2+}]i$).

**RNA sequencing sample preparation and Ingenuity Pathway analysis**

RNA was extracted using small [<200 nt] and large [>200 nt] nucleotide separation from BZ and RZ myocardial tissue from control *Runx1*$^{fl/fl}$ and *Runx1*$^{\Delta/\Delta}$ mice pre-MI and 1 day post-MI as well as whole LV myocardial tissue from *CBFβ*$^{fl/fl}$ and *CBFβ*$^{\Delta/\Delta}$ pre-MI and 7 days post-MI. The raw fastq files containing single-end 1 × 75 bp reads were pre-processed with Cutadapt (v.1.8) and Sickle (v.0.940) software (https://github.com/najoshi/sickle) to remove the 3' end adaptor and to trim the very low-quality reads, respectively. The quality threshold was set to 10 and no reads shorter than 54 bp were allowed to remain (sickle flags: -q 10, -l 54). The pre-processed reads were then aligned to the reference genome (Ensembl GRCm38.95) with Hisat264 (v2.1.0) and transcript expression quantification was performed using StringTie65 (v1.3.5). Gene level count matrices were generated with prepDE.py script as instructed in StringTie manual (http://www.ccb.jhu.edu/software/stringtie/index.shtml?t=manual) and differential expression were analysed using edgeR66 R-package to provide statistically significant gene-lists for a variety of between-groups comparisons. Gene lists and expression values were uploaded to Ingenuity® Pathway Analysis (IPA) software (QIAGEN Inc.)[2]. The core analysis feature was used to interpret the differentially expressed data (FDR ≤ 0.05), including biological (canonical) pathway analysis and an activity analysis z-score giving a pathway prediction of inhibition (blue) or activation (orange). Comparison analysis between specific groups were carried out to visualise relevant canonical pathways using right-tailed Fisher's Exact Test and corrected for multiple testing using Benjamini Hochberg.

**Electron Microscopy**

Hearts were perfused with cardioplegic solution, immediately followed by perfusion-fixation with iso-osmotic Karnovsky's fixative (2.4% sodium cacodylate, 0.75% paraformaldehyde, 0.75% glutaraldehyde). BZ and remote LV regions were then washed with 0.1 M sodium cacodylate, post-fixed in 1% $OsO_4$ for 1 h, dehydrated in graded acetone, and embedded in Epon-Araldite resin as described before[3]. All sections were placed on formvar-coated copper/palladium slot-grids, post stained with 2% aqueous uranyl acetate and Reynold's lead citrate. Colloidal gold particles (15 nm) were added to both surfaces of the thick sections to serve as fiducial markers for tilt series alignment. For tomographic imaging, the specimen holder was tilted from +60° to –60° at 1° intervals. For dual-axis tilt series the specimen was then rotated by 90° in the X-Y plane, and another +60° to –60° tilt series was taken. The images from each tilt-series were aligned by fiducial marker tracking and back-projected to generate two single full-thickness reconstructed volumes (tomograms), which were then combined to generate a single high-resolution 3D reconstruction of the original partial cell volume. Mitochondrial density was calculated in each cardiomyocyte, represented as a percentage of the total cellular area. The number of mitochondria were also counted in the same cells, with mitochondrial size determined by dividing the total mitochondrial area by the number of mitochondria. Mitochondria were classified as damaged if: cristae and outer membrane integrity was disrupted (dissolved/damaged cristae); evidence of fusion of mitochondria with autophagocytic vesicles was present; or if there were mitophagosomes present. Damaged mitochondria were counted in each cell and the proportion present calculated by dividing the number of damaged mitochondria by the total.

**RNA isolation, cDNA synthesis and real-time qPCR analysis**

Total RNA was extracted from cardiomyocytes from the LV using the miRNeasy Mini Kit (Qiagen, UK) with DNase I treatment (Qiagen, UK). 1 µg of RNA was then reverse transcribed to cDNA with Omniscript reverse transcriptase (Omniscipt Reverse Transcription kit, Qiagen, UK). Real-time quantitative PCR (qPCR) reactions were run using cDNA with a Runx1 Taqman Gene Expression Assay (Amplicon length 81bp, Mm01213404_m1) and Taqman Universal Mastermix Mix II, no UNG (both ThermoFisher Scientific, UK) in a final volume of 10 µL with the following cycling conditions: hold for 2 min at 50°C followed by 10 min at 95°C then 40 cycles of 95°C for 10 min, 60°C for 1 min. Relative mRNA levels were analysed using comparative Ct calculations ($2^{-\Delta Ct}$); normalised to peptidylprolyl isomerase B; *Ppib* (ThermoFisher Scientific, UK).

**Adenoviral knockdown of Runx1 in the border zone region**

*Runx1* shRNA adenovirus encoded *Runx1* shRNA[4] short hairpin forward 5'-GATCCCCGGGCCCTCCTACCATCTATACTACTCGAGTAGTATAGATGGTAGGAGGGCTTTTTGG-3' and reverse 5'-AATTCCAAAAAGCCCTCCTACCATCTATACTACTCGAGTAGTATAGATGGTAGGAGGGCCCGGG-3'. Scramble shRNA short hairpin contained forward: 5'-GATCCCCGGCAACAAGATGAAGAGCACCAACTCGAGTTGGTGCTCTTCATCTTGTTGTTTTTG-3' and reverse 5'-AATTCAAAAACAACAAGATGAAGAGCACCAACTCGAGTTGGTGCTCTTCATCTTGTTGCCGGG-3'. Ad-scramble-shRNA and Ad-*Runx1*-shRNA were validated *in vitro* using the mouse endothelial IP1B cell line (obtained from the American Type Culture Collection; ATCC) (Extended Figure 3).

**Generation of AAV-scramble-shRNA and AAV-Runx1-shRNA**

shRNA sequences targeting murine *Runx1* were generated using the short hairpin sequences in the adenoviral vectors for both *Runx1* and scramble. Briefly, the complementary oligos of shRNA (Eurofins genomics, Germany) were annealed and cloned into a pZac2.1 vector, which was then used to produced recombinant AAV vectors in the AAV Vector Unit at IGGEB Triest, as described previously[5]. Viral stocks were obtained by PEG precipitation and two subsequent $CsCl_2$ Gradient centrifugations. Titration of AAV viral particles was performed by real-time PCR quantification of the number of packaged viral genomes as described previously[6].

References

1. McCarroll CS, He W, Foote K, Bradley A, McGlynn K, Vidler F, Nixon C, Nather K, Fattah C, Riddell A, Bowman P, Elliott EB, Bell M, Hawksby C, MacKenzie SM, Morrison LJ, Terry A, Blyth K, Smith GL, McBride MW, Kubin T, Braun T, Nicklin SA, Cameron ER and Loughrey CM. Runx1 Deficiency Protects Against Adverse Cardiac Remodeling After Myocardial Infarction. *Circulation*. 2018;137:57-70.

2. Kramer A, Green J, Pollard Jr J, Tugendreich S. Causal analysis approaches in Ingenuity Pathway Analysis. Bioinformatics. 2014;30(4):523030.

3. Rog-Zielinska EA, Jonhston CM, O'Toole ET, Morphew M, Hoenger A, Kohl P. Electron tomography of rabbit cardiomyocyte three-dimensional ultrastructure. Prog Biophys Mol Biol. 2016;121(2):77-84.

4. Wang J, Wang X, Holz JD, Rutkowski T, Wang Y, Zhu Z and Dong Y. Runx1 is critical for PTH-induced onset of mesenchymal progenitor cell chondrogenic differentiation. *PloS one*. 2013;8:e74255.

5. Ayuso E, Blouin V, Lock M, McGorray S, Leon X, Alvira MR, Auricchio A, Bucher S, Chtarto A, Clark KR, Darmon C, Doria M, Fountain W, Gao G, Gao K, Giacca M, Kleinschmidt J, Leuchs B, Melas C, Mizukami H, Muller M, Noordman Y, Bockstael O, Ozawa K, Pythoud C, Sumaroka M, Surosky R, Tenenbaum L, van der Linden I, Weins B, Wright JF, Zhang X, Zentilin L, Bosch F, Snyder RO and Moullier P. Manufacturing and characterization of a recombinant adeno-associated virus type 8 reference standard material. *Hum Gene Ther*. 2014;25:977-87.

6. Arsic N, Zacchigna S, Zentilin L, Ramirez-Correa G, Pattarini L, Salvi A, Sinagra G and Giacca M. Vascular endothelial growth factor stimulates skeletal muscle regeneration in vivo. *Mol Ther*. 2004;10:844-54.

### Extended Figure 1

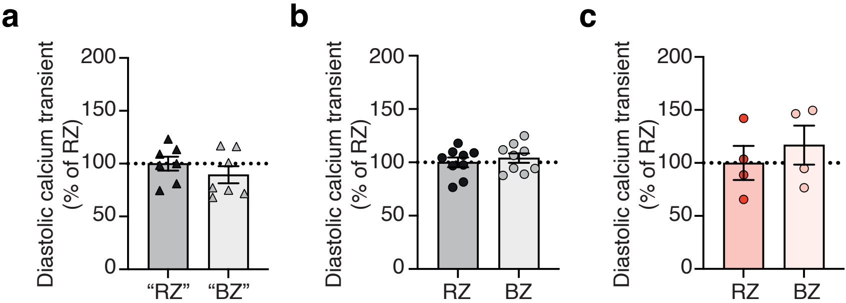

**Extended Figure 2**

Border zone *vs.* Remote zone

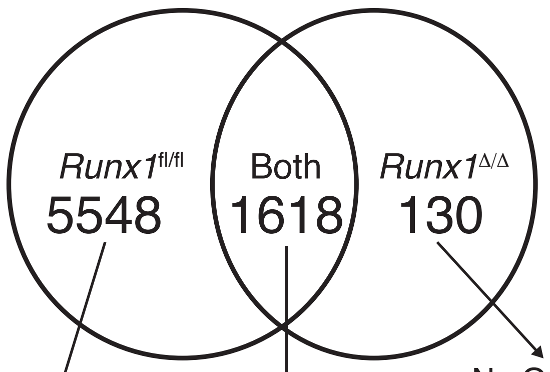


No OxPhos genes differentially regulated

No OxPhos genes differentially regulated

Day 7 Post-MI *vs.* Pre-MI

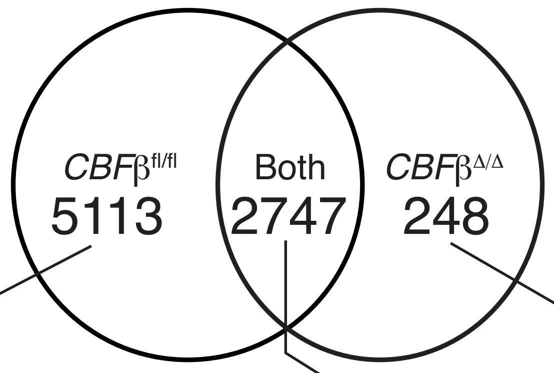


No OxPhos genes differentially regulated

##### *Runx1*$^{fl/fl}$

| | FDR | Log Ratio | Intensity |
|---|---|---|---|
| ATP5A1 | 0.0039523 | -0.76 | 11.5 |
| ATP5B | 0.0022955 | -0.67 | 11.9 |
| ATP5C1 | 0.0116387 | -0.63 | 9.5 |
| ATP5D | 0.0058572 | -0.47 | 8.6 |
| ATP5F1 | 0.0092230 | -0.63 | 9.0 |
| ATP5G2 | 0.0480540 | -0.25 | 6.6 |
| ATP5G3 | 0.0029577 | -0.72 | 9.5 |
| ATP5H | 0.0359252 | -0.46 | 9.1 |
| ATP5J2 | 0.0474604 | -0.40 | 8.6 |
| ATP5L | 0.0105228 | -0.52 | 7.6 |
| ATP5O | 0.0073731 | -0.45 | 8.9 |
| COX1 | 0.0016757 | -0.80 | 16.4 |
| COX2 | 0.0010174 | -0.65 | 13.9 |
| COX3 | 0.0010280 | -0.75 | 14.2 |
| COX4I1 | 0.0004623 | -0.62 | 9.3 |
| COX5A | 0.0020023 | -0.64 | 9.1 |
| COX6A1 | 0.0038411 | 0.33 | 5.8 |
| COX6A2 | 0.0129188 | -0.52 | 9.9 |
| COX6C | 0.0235638 | -0.47 | 5.4 |
| COX7A1 | 0.0191882 | -0.60 | 8.6 |
| COX7A2 | 0.0229588 | -0.41 | 7.6 |
| COX7B | 0.0305723 | -0.47 | 8.9 |
| CYTB | 0.0122457 | -0.63 | 15.3 |
| ND1 | 0.0003733 | -0.83 | 14.4 |
| ND2 | 0.0022125 | -0.73 | 13.9 |
| ND3 | 0.0017862 | -0.57 | 10.1 |
| ND4 | 0.0022339 | -0.71 | 14.4 |
| ND4L | 0.0031282 | -0.87 | 11.1 |
| ND5 | 0.0052253 | -0.80 | 14.3 |
| NDUFA1 | 0.0231656 | -0.43 | 7.0 |
| NDUFA10 | 0.0032834 | -0.69 | 8.5 |
| NDUFA11 | 0.0466337 | -0.37 | 5.9 |
| NDUFA13 | 0.0229270 | -0.39 | 8.2 |
| NDUFA2 | 0.0381739 | -0.29 | 7.1 |
| NDUFA4 | 0.0062586 | -0.47 | 8.5 |
| NDUFA5 | 0.0127572 | -0.63 | 7.8 |
| NDUFA6 | 0.0378071 | -0.44 | 6.6 |
| NDUFA7 | 0.0418919 | -0.27 | 6.9 |
| NDUFA8 | 0.0016706 | -0.66 | 7.6 |
| NDUFA9 | 0.0110475 | -0.59 | 8.6 |
| NDUFAB1 | 0.0045633 | -0.71 | 3.6 |
| NDUFB10 | 0.0243309 | -0.65 | 7.7 |
| NDUFB11 | 0.0338063 | -0.44 | 7.8 |
| NDUFB2 | 0.0443747 | -0.39 | 6.7 |
| NDUFB3 | 0.0319472 | -0.42 | 6.6 |
| NDUFB4 | 0.0214261 | -0.47 | 4.1 |
| NDUFB5 | 0.0196200 | -0.53 | 7.4 |
| NDUFB6 | 0.0082544 | -0.48 | 6.6 |
| NDUFB7 | 0.0174701 | -0.36 | 7.5 |
| NDUFB8 | 0.0382447 | -0.41 | 7.9 |
| NDUFB9 | 0.0063922 | -0.55 | 8.6 |
| NDUFS1 | 0.0090476 | -0.75 | 9.2 |
| NDUFS2 | 0.0041991 | -0.65 | 9.4 |
| NDUFS3 | 0.0070739 | -0.66 | 7.7 |
| NDUFS4 | 0.0103826 | -0.58 | 6.8 |
| NDUFS6 | 0.0208048 | -0.46 | 7.5 |
| NDUFS7 | 0.0002880 | -0.74 | 7.8 |
| NDUFS8 | 0.0159283 | -0.40 | 7.2 |
| NDUFV1 | 0.0013644 | -0.68 | 8.5 |
| NDUFV2 | 0.0130075 | -0.53 | 8.3 |
| NDUFV3 | 0.0413884 | -0.39 | 7.5 |
| SDHA | 0.0034819 | -0.77 | 10.2 |
| SDHB | 0.0107764 | -0.57 | 9.2 |
| SDHC | 0.0005796 | -0.71 | 8.1 |
| SDHD | 0.0120490 | -0.66 | 8.4 |
| UQCR10 | 0.0327368 | -0.41 | 7.7 |
| UQCR11 | 0.0151261 | -0.40 | 7.4 |
| UQCRB | 0.0318067 | -0.46 | 8.2 |
| UQCRC1 | 0.0011091 | -0.72 | 9.4 |
| UQCRC2 | 0.0157486 | -0.61 | 9.4 |
| UQCRFS1 | 0.0055431 | -0.67 | 9.2 |
| UQCRQ | 0.0277681 | -0.36 | 8.1 |

##### *CBF*β$^{fl/fl}$

| | FDR | Log Ratio | Intensity |
|---|---|---|---|
| ATP5D | 0.0000073 | -0.43 | 8.7 |
| ATP5G1 | 0.0000022 | -0.59 | 9.3 |
| ATP5G2 | 0.0000072 | -0.45 | 6.8 |
| ATP5G3 | 0.0000122 | -0.59 | 9.9 |
| ATP5J | 0.0000055 | -0.77 | 8.6 |
| ATP5J2 | 0.0000322 | -0.65 | 8.9 |
| ATP5L | 0.0121140 | -0.72 | 7.8 |
| COX1 | 0.0002330 | -0.75 | 16.4 |
| COX2 | 0.0007620 | -0.70 | 13.9 |
| COX3 | 0.0024200 | -0.77 | 14.0 |
| COX4I1 | 0.0018550 | -0.44 | 9.5 |
| COX6A2 | 0.0000268 | -0.61 | 10.0 |
| COX7A2 | 0.0000002 | -0.61 | 8.2 |
| CYTB | 0.0001400 | -0.75 | 15.1 |
| ND1 | 0.0028610 | -0.75 | 14.2 |
| ND2 | 0.0003630 | -0.80 | 13.7 |
| ND3 | 0.0200290 | -0.63 | 10.1 |
| ND4 | 0.0004380 | -0.77 | 14.3 |
| ND4L | 0.0005770 | -0.90 | 10.7 |
| ND5 | 0.0003310 | -0.83 | 14.0 |
| NDUFA11 | 0.0338320 | -0.51 | 0.0 |
| NDUFA12 | 0.0000033 | -0.68 | 7.6 |
| NDUFA13 | 0.0000001 | -0.66 | 8.4 |
| NDUFA2 | 0.0000064 | -0.54 | 7.4 |
| NDUFA4 | 0.0000604 | -0.69 | 8.9 |
| NDUFA5 | 0.0000006 | -0.72 | 8.0 |
| NDUFA6 | 0.0000006 | -0.61 | 6.8 |
| NDUFA7 | 0.0000249 | -0.58 | 7.0 |
| NDUFA8 | 0.0000004 | -0.53 | 8.1 |
| NDUFAB1 | 0.0000059 | -0.69 | 3.7 |
| NDUFB10 | 0.0010650 | -0.68 | 8.1 |
| NDUFB6 | 0.0000001 | -0.55 | 6.9 |
| NDUFS4 | 0.0091320 | -0.44 | 7.2 |
| NDUFS7 | 0.0000022 | -0.56 | 8.1 |
| NDUFV3 | 0.0000485 | -0.60 | 7.6 |
| UQCR11 | 0.0000039 | -0.68 | 7.6 |

##### *CBF*β$^{fl/fl}$ and *CBF*β$^{Δ/Δ}$

| *CBF*β$^{fl/fl}$ FDR | Log Ratio | Intensity | | *CBF*β$^{Δ/Δ}$ FDR | Log Ratio | Intensity |
|---|---|---|---|---|---|---|
| 0.00000009 | -0.73 | 11.7 | ATP5A1 | 0.0146323 | -0.29 | 11.7 |
| 0.00000003 | -0.69 | 12.0 | ATP5B | 0.0054477 | -0.29 | 12.0 |
| 0.00000005 | -0.71 | 9.7 | ATP5C1 | 0.0306031 | -0.23 | 9.7 |
| 0.00000028 | -0.70 | 9.3 | ATP5F1 | 0.0437999 | -0.24 | 9.3 |
| 0.00000043 | -0.59 | 9.4 | ATP5H | 0.0183580 | -0.24 | 9.4 |
| 0.00000491 | -0.62 | 9.1 | ATP5O | 0.0321916 | -0.28 | 9.1 |
| 0.00000046 | -0.60 | 9.4 | COX5A | 0.0360319 | -0.22 | 9.4 |
| 0.00000127 | -0.53 | 8.7 | COX6B1 | 0.0035017 | -0.30 | 8.7 |
| 0.00000068 | -0.71 | 5.5 | COX6C | 0.0215390 | -0.30 | 5.5 |
| 0.00000002 | -1.11 | 8.9 | COX7A1 | 0.0017462 | -0.52 | 8.9 |
| 0.00000013 | -0.69 | 9.1 | COX7B | 0.0411036 | -0.23 | 9.1 |
| 0.01793716 | -0.12 | 6.7 | COX8A | 0.0492167 | -0.11 | 6.7 |
| 0.00000002 | -0.72 | 7.1 | NDUFA1 | 0.0039720 | -0.31 | 7.1 |
| 0.00000036 | -0.60 | 8.6 | NDUFA10 | 0.0072908 | -0.28 | 8.6 |
| 0.00000184 | -0.68 | 7.3 | NDUFA3 | 0.0353796 | -0.28 | 7.3 |
| 0.00000002 | -0.73 | 8.8 | NDUFA9 | 0.0016777 | -0.35 | 8.8 |
| 0.00002809 | -0.66 | 7.2 | NDUFB1 | 0.0161446 | -0.38 | 7.2 |
| 0.00000002 | -0.67 | 8.2 | NDUFB11 | 0.0033537 | -0.29 | 8.2 |
| 0.00000399 | -0.61 | 7.0 | NDUFB2 | 0.0387330 | -0.26 | 7.0 |
| 0.00000344 | -0.50 | 7.0 | NDUFB3 | 0.0112614 | -0.26 | 7.0 |
| 0.00000002 | -0.74 | 4.3 | NDUFB4 | 0.0205777 | -0.26 | 6.9 |
| 0.00000002 | -0.64 | 7.7 | NDUFB5 | 0.0085608 | -0.25 | 7.7 |
| 0.00000041 | -0.52 | 7.8 | NDUFB7 | 0.0408524 | -0.19 | 7.8 |
| 0.00000031 | -0.63 | 8.3 | NDUFB8 | 0.0061780 | -0.30 | 8.3 |
| 0.00000005 | -0.67 | 8.9 | NDUFB9 | 0.0421062 | -0.21 | 8.9 |
| 0.00000001 | -0.83 | 9.5 | NDUFS1 | 0.0008842 | -0.41 | 9.5 |
| 0.00000004 | -0.67 | 9.6 | NDUFS2 | 0.0078584 | -0.27 | 9.6 |
| 0.00000065 | -0.68 | 8.0 | NDUFS3 | 0.0150749 | -0.30 | 8.0 |
| 0.00000020 | -0.70 | 7.9 | NDUFS6 | 0.0221499 | -0.27 | 7.9 |
| 0.00000014 | -0.65 | 7.5 | NDUFS8 | 0.0121548 | -0.27 | 7.5 |
| 0.00000064 | -0.65 | 8.6 | NDUFV1 | 0.0322420 | -0.25 | 8.6 |
| 0.00000074 | -0.66 | 8.7 | NDUFV2 | 0.0382739 | -0.25 | 8.7 |
| 0.00000018 | -0.87 | 10.4 | SDHA | 0.0149728 | -0.36 | 10.4 |
| 0.00000010 | -0.69 | 9.4 | SDHB | 0.0162599 | -0.27 | 9.4 |
| 0.00000019 | -0.67 | 8.4 | SDHC | 0.0200184 | -0.26 | 8.4 |
| 0.00000007 | -0.64 | 8.8 | SDHD | 0.0144046 | -0.25 | 8.8 |
| 0.00000073 | -0.45 | 8.0 | UQCR10 | 0.0171973 | -0.20 | 8.0 |
| 0.00000003 | -0.69 | 8.5 | UQCRB | 0.0079927 | -0.27 | 8.5 |
| 0.00000008 | -0.66 | 9.7 | UQCRC1 | 0.0461082 | -0.21 | 9.7 |
| 0.00000004 | -0.65 | 9.6 | UQCRC2 | 0.0029839 | -0.30 | 9.6 |
| 0.00000014 | -0.72 | 9.5 | UQCRFS1 | 0.0089345 | -0.31 | 9.5 |
| 0.00005959 | -0.58 | 2.1 | UQCRH | 0.0106783 | -0.38 | 2.1 |
| 0.00000048 | -0.66 | 8.4 | UQCRQ | 0.0277046 | -0.26 | 8.4 |

Downregulated

Upregulated

#### Extended Figure 3

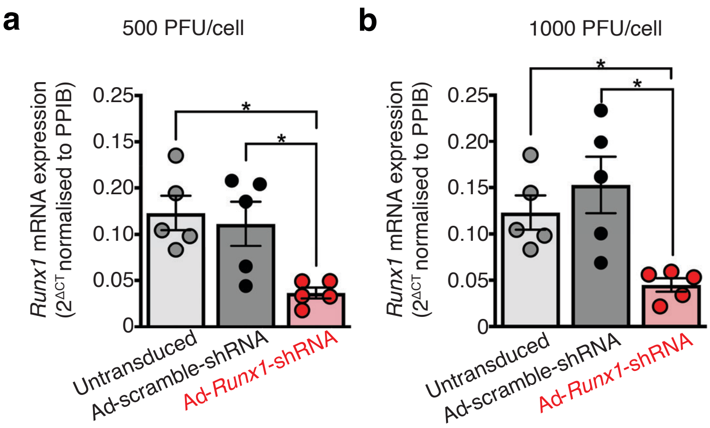

### Extended Figure 4

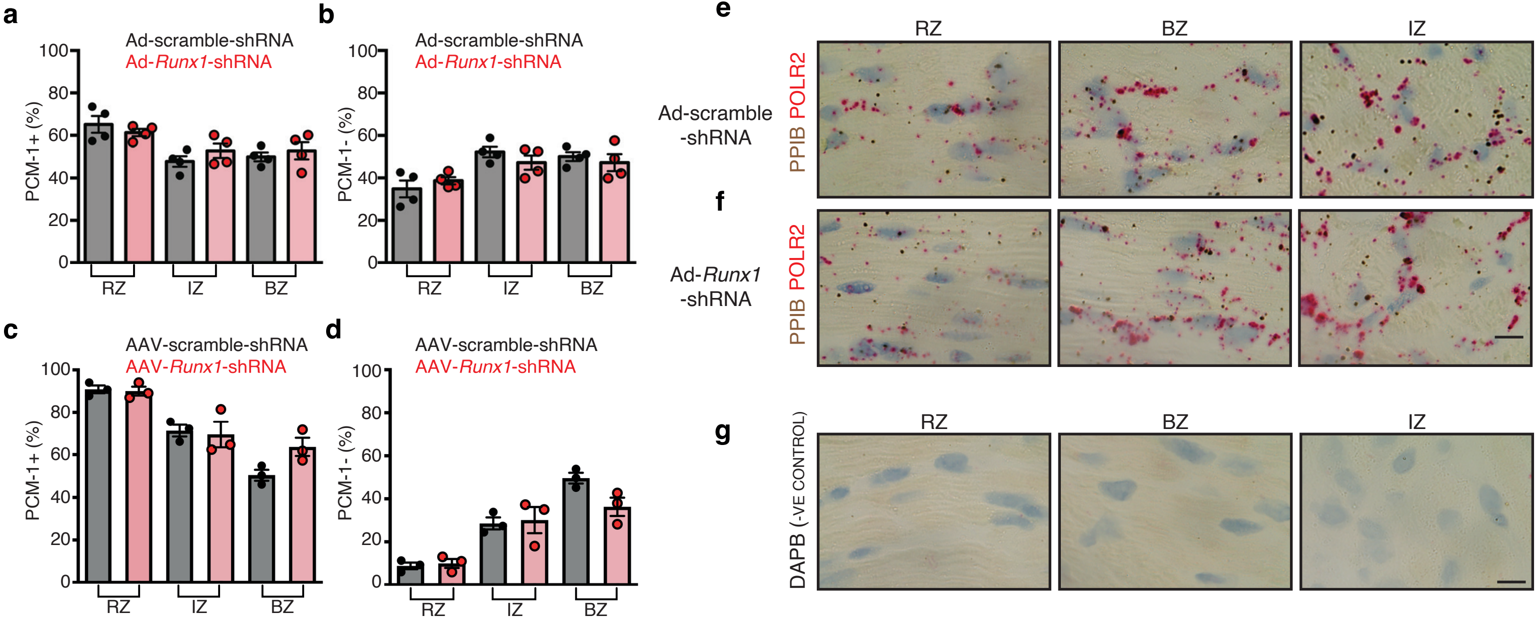

**Extended Table 1 - Calcium**

| | pre-MI C57BL6 | | | | Day 1 Post-MI C57BL6 | | | | Day 1 Post-MI Runx1 deficient | | | | Day 1 Post-MI Runx1 floxed | | | |
|---|---|---|---|---|---|---|---|---|---|---|---|---|---|---|---|---|
| | RZ | | BZ | | RZ | | BZ | | RZ | | BZ | | RZ | | BZ | |
| | **MEAN** | *SEM* | **MEAN** | SEM | **MEAN** | *SEM* | **MEAN** | *SEM* | **MEAN** | *SEM* | **MEAN** | SEM | **MEAN** | *SEM* | **MEAN** | *SEM* |
| CALCIUM TRANSIENT PEAK (nmol.L$^{-1}$) | **405.53** | *19.35* | **406.01** | *18.51* | **418.44** | *21.41* | **263.90*** | *19.30* | **317.28** | *17.61* | **373.15** | *67.08* | **540.79** | *40.07* | **313.85*** | *27.34* |
| CALCIUM TRANSIENT MINIMUM (nmol.L$^{-1}$) | **166.46** | *7.50* | **171.96** | *8.04* | **166.38** | *10.92* | **144.35** | *6.66* | **131.96** | *21.18* | **164.08** | *46.72* | **218.32** | *18.17* | **173.90** | *10.10* |
| CALCIUM TRANSIENT AMPLITUDE (nmol.L$^{-1}$) | **239.07** | *16.12* | **234.05** | *16.82* | **252.06** | *19.47* | **119.54*** | *16.92* | **185.26** | *6.24* | **209.07** | *27.10* | **322.47** | *24.51* | **139.95*** | *19.56* |
| CAFFEINE-INDUCED CALCIUM TRANSIENT AMPLITUDE (nmol.L$^{-1}$) | **652.40** | *14.51* | **663.28** | *30.62* | **606.65** | *27.49* | **431.51*** | *43.18* | **457.62** | *82.25* | **474.42** | *79.50* | **413.29** | *18.66* | **315.88*** | *28.84* |
| SERCA ACTIVITY (KSERCA (s$^{-1}$)) | **10.35** | *0.77* | **9.11** | *0.96* | **9.62** | *1.21* | **4.51*** | *0.76* | **6.16** | *0.68* | **7.53** | *2.02* | **6.56** | *0.57* | **4.63*** | *0.40* |
| TIME CONSTANT OF CAFFEINE INDUCED TRANSIENT DECAY (NCX ACTIVITY (s$^{-1}$)) | **1.16** | *0.10* | **0.98** | *0.08* | **1.01** | *0.21* | **1.24** | *0.39* | **0.72** | *0.14* | **0.72** | *0.13* | **0.41** | *0.02* | **0.41** | *0.02* |
| CALCIUM TRANSIENT PEAK TO MINIMUM RATIO | **2.45** | *0.12* | **2.38** | *0.11* | **2.56** | *0.16* | **1.84*** | *0.12* | **2.53** | *0.26* | **2.57** | *0.40* | **2.49** | *0.08* | **1.80*** | *0.09* |

**Extended Table 2 - Echocardiography**

| | Ad-scramble-shRNA | | | | | Ad-Runx1-shRNA | | | | |
|---|---|---|---|---|---|---|---|---|---|---|
| | Fractional shortening (%) | LVIDd (mm) | LVIDs (mm) | LVPWd (mm) | LVPWs (mm) | Fractional shortening (%) | LVIDd (mm) | LVIDs (mm) | LVPWd (mm) | LVPWs (mm) |
| Pre-MI | 42.74 | 3.80 | 2.18 | 1.24 | 1.53 | 43.81 | 3.81 | 2.14 | 1.27 | 1.58 |
| 1-day post-MI | 26.43 | 4.12 | 3.03 | 1.14 | 1.27 | 34.73 | 3.68 | 2.41 | 1.20 | 1.38 |
| 2 days post-MI | 26.38 | 4.26 | 3.13 | 1.06 | 1.32 | 38.32 | 3.71 | 2.29 | 1.11 | 1.38 |
| 7 days post-MI | 28.46 | 3.92 | 2.81 | 1.38 | 1.54 | 40.88 | 3.95 | 2.33 | 1.22 | 1.54 |

| | AAV-scramble-shRNA | | | | | AAV-Runx1-shRNA | | | | |
|---|---|---|---|---|---|---|---|---|---|---|
| | Fractional shortening (%) | LVIDd (mm) | LVIDs (mm) | LVPWd (mm) | LVPWs (mm) | Fractional shortening (%) | LVIDd (mm) | LVIDs (mm) | LVPWd (mm) | LVPWs (mm) |
| Pre-MI | 46.58 | 3.55 | 1.91 | 1.31 | 1.80 | 45.92 | 3.61 | 1.96 | 1.19 | 1.65 |
| 1-day post-MI | 27.62 | 4.02 | 2.93 | 1.04 | 1.28 | 37.66 | 3.89 | 2.43 | 1.20 | 1.54 |
| 7 days post-MI | 28.21 | 4.30 | 3.10 | 1.26 | 1.51 | 37.21 | 3.92 | 2.48 | 1.41 | 1.71 |

| | PROTOCOL 1 | | | | | | | | | |
|---|---|---|---|---|---|---|---|---|---|---|
| | Vehicle | | | | | Ro5-3335 | | | | |
| | Fractional shortening (%) | LVIDd (mm) | LVIDs (mm) | LVPWd (mm) | LVPWs (mm) | Fractional shortening (%) | LVIDd (mm) | LVIDs (mm) | LVPWd (mm) | LVPWs (mm) |
| Pre-MI | 41.45 | 3.46 | 2.03 | 1.36 | 1.59 | 42.72 | 3.44 | 1.97 | 1.29 | 1.72 |
| 1-day post-MI | 30.14 | 3.83 | 2.68 | 1.25 | 1.48 | 37.60 | 3.66 | 2.31 | 1.15 | 1.53 |
| 3 days post-MI | 32.75 | 3.35 | 2.26 | 1.50 | 1.85 | 40.35 | 3.71 | 2.26 | 1.42 | 1.77 |
| 7 days post-MI | 30.13 | 3.56 | 2.49 | 1.33 | 1.52 | 38.77 | 3.73 | 2.29 | 1.36 | 1.61 |

| | PROTOCOL 2 | | | | | | | | | |
|---|---|---|---|---|---|---|---|---|---|---|
| | Vehicle | | | | | Ro5-3335 | | | | |
| | Fractional shortening (%) | LVIDd (mm) | LVIDs (mm) | LVPWd (mm) | LVPWs (mm) | Fractional shortening (%) | LVIDd (mm) | LVIDs (mm) | LVPWd (mm) | LVPWs (mm) |
| Pre-MI | 41.83 | 3.29 | 1.92 | 1.33 | 1.67 | 44.80 | 3.32 | 1.83 | 1.25 | 1.64 |
| 1-day post-MI | 26.23 | 3.82 | 2.83 | 1.21 | 1.35 | 32.20 | 3.73 | 2.54 | 1.10 | 1.46 |
| 3 days post-MI | 24.58 | 3.91 | 2.96 | 0.94 | 1.24 | 38.64 | 3.49 | 2.17 | 1.28 | 1.78 |
| 7 days post-MI | 26.60 | 4.09 | 3.01 | 0.93 | 1.15 | 33.35 | 3.77 | 2.52 | 1.24 | 1.74 |

| | $CBF\beta^{fl/fl}$ | | | | | $CBF\beta^{\Delta/\Delta}$ | | | | |
|---|---|---|---|---|---|---|---|---|---|---|
| | Fractional shortening (%) | LVIDd (mm) | LVIDs (mm) | LVPWd (mm) | LVPWs (mm) | Fractional shortening (%) | LVIDd (mm) | LVIDs (mm) | LVPWd (mm) | LVPWs (mm) |
| Pre-MI | 41.77 | 3.16 | 1.82 | 1.46 | 1.71 | 42.72 | 3.44 | 1.97 | 1.29 | 1.72 |
| 1-day post-MI | 30.16 | 3.83 | 2.68 | 1.25 | 1.48 | 37.60 | 3.66 | 2.31 | 1.15 | 1.53 |
| 3 days post-MI | 32.75 | 3.35 | 2.26 | 1.50 | 1.85 | 40.35 | 3.71 | 2.26 | 1.42 | 1.77 |
| 7 days post-MI | 30.13 | 3.56 | 2.49 | 1.33 | 1.52 | 38.77 | 3.73 | 2.29 | 1.36 | 1.61 |
